# Early MATR3 loss and distinct neurodegenerative molecular signatures precede the onset of neuropathology in motor neurons and Purkinje cells of MATR3 S85C knock-in mouse model of ALS

**DOI:** 10.64898/2026.08.26.747343

**Authors:** Katarina Maksimovic, Rambabu Majji, Jhune Rizsan Santos, Cadia Chan, Anneka Zelaya, Jooyun Lee, Michelle Dias, Oxana B. Gluscencova, Mohieldin M. M. Youssef, Sandra Kim, Tiara Noronha, Christine Lai, Yiting Fan, Mark N. Metri, Justin You, Ching Serena Kao, Lu-Yang Wang, Julie L. Lefebvre, Michael D. Wilson, Hari Krishna Yalamanchili, Jeehye Park

## Abstract

Amyotrophic lateral sclerosis (ALS) is a motor neuron disease, leading to progressive muscle weakness and motor impairment. Growing evidence indicates that cerebellar Purkinje cells, which play a central role in motor coordination, are also affected in ALS. However, it is unclear whether the molecular events that initiate neurodegeneration in these ALS-relevant motor-controlling neurons are shared or distinct. Here, we used a MATR3 S85C knock-in (KI) mouse model of early-stage ALS with stage-specific motor phenotypes and selective vulnerability of motor neurons and Purkinje cells to decipher the molecular events underlying neurodegeneration in these two neuronal populations. We found that a profound reduction in detectable MATR3 S85C immunoreactivity (hereafter referred to as MATR3 loss) in both motor neurons and Purkinje cells precedes the onset of motor dysfunction and neuropathology, implicating MATR3 loss as the earliest detectable molecular event. Our bulk cerebellar RNA profiling and motor neuron-specific RNA profiling data at the onset of MATR3 loss revealed distinct molecular signatures. In the cerebellum, *Ngfr* expression emerged in Purkinje cells before the onset of neuronal loss and remained elevated throughout the disease course. This increase was accompanied by activation of the JNK-mediated cell death pathway. In the motor neurons, elevated *Fgf21* and integrated stress response (ISR) gene expression were the first to be observed and persisted throughout disease progression, consistent with previous findings in SOD1 mouse models. Our findings provide mechanistic insights into the initiation of neurodegeneration in ALS-relevant motor-controlling neurons and implicate potential neuron type-specific targets for future therapeutics.

## INTRODUCTION

Amyotrophic lateral sclerosis (ALS) is a movement disorder associated with muscle atrophy and paralysis^1, 2^. Since its pathological description by Jean-Martin Charcot about 150 years ago, ALS has been historically defined by motor neuron loss^3^. The discovery of ALS-causing genes and subsequent studies of their functions as well as transcriptome profiling studies in spinal cord and motor neurons have provided significant insights into the mechanisms underlying motor neuron degeneration^4-14^. Despite extensive research, the mechanisms that initiate motor neuron degeneration and drive disease progression remain unclear. Understanding how neurodegeneration is initiated and progresses would be critical for the development of early diagnostic tools and preventative therapeutic strategies.

Several groups have performed longitudinal, motor neuron-specific transcriptional profiling in SOD1 G93A, G37R, or G85R mouse models to define early, progressive molecular changes within motor neurons during disease development^10-14^. In particular, selective profiling of spinal motor neurons through laser capture microdissection or translating ribosome affinity purification (TRAP)-based approaches has identified convergent transcriptional signatures involving complement, ER stress, and regenerative/injury responses that emerge before symptom onset and persist with disease progression across SOD1 G93A, G37R, and G85R models^11,12,14^. However, these transgenic models have been generated by introducing high copy numbers of the mutant human SOD1 transgene, so it remains unclear whether the observed transcriptional responses result from the overload of the protein or are specifically driven by the disease-linked mutation. Furthermore, little is known when the disease-causing mutation begins to alter protein properties and function and initiate the molecular events that ultimately lead to neurodegeneration.

Although ALS is classically associated with motor neuron pathology, there is growing evidence revealing pathological changes in the cerebellum and Purkinje cells, which control balance, motor coordination, and gait refinement^15^. Neuroimaging studies on sporadic and familial ALS cases presented notable cerebellar atrophy and gray matter loss^16-23^, suggesting that cerebellar pathology contributes to gait impairment. Consistently, neuropathological studies on postmortem cerebellar tissue from ALS patients and genetic ALS animal models including SOD1 transgenic mice have revealed Purkinje cell loss^24-35^. In addition, ubiquitin, p62 (sequestosome-1) and/or TDP-43 (TAR DNA-binding protein 43)-positive inclusions were observed in the cerebellar tissue in ALS patients^36-42^. Furthermore, recent transcriptomic profiling studies revealed extensive molecular alterations in the cerebella of ALS patient samples^43-46^, supporting the idea that cerebellar pathology may contribute to ALS pathogenesis. However, the mechanisms underlying cerebellar pathology and Purkinje cell degeneration in ALS remain poorly understood, and it is unclear whether the same molecular pathways that underlie motor neuron degeneration contribute to cerebellar and Purkinje cell degeneration.

Mutations in *MATR3*, which encodes for a ubiquitously expressed protein with roles in DNA- and RNA-associated functions, have been linked to ALS^47-49^. Particularly, the serine 85 to cysteine mutation (or S85C) is the most common familial mutation in MATR3 associated with a slow progressive form of ALS^47^. We previously generated a novel ALS mouse model, MATR3 S85C knock-in (KI) mice, that closely represent the natural disease progression^35^. This mouse line harbors a single S85C missense mutation in endogenous mouse *Matr3* gene, thus expressing the mutant protein, MATR3 S85C, at the physiological level^35^. We found that these KI mice exhibit gradual progression of motor dysfunction and hindlimb weakness. In addition, the mutant mice exhibit loss of Purkinje cells in the cerebellum and neuromuscular junction (NMJ) defects^35^, which is an early pathological feature of ALS, occurring long before the loss of motor neuron cell bodies in the spinal cord^50-54^. Notably, our neuropathological data showed a striking loss of nuclear MATR3 S85C immunoreactivity (or MATR3 loss) of the affected neuronal types including alpha motor neurons and Purkinje cells, but not in neighboring neurons^35^, indicating that specific cell types are impacted by the ALS-linked S85C mutation. Given that MATR3 S85C KI mice exhibit slow progressive ALS-like phenotypes together with the disease-relevant, cell type-specific impact of the mutation, this model offers enhanced disease relevance and provides an opportunity to dissect the temporal emergence and mechanisms underlying early disease pathogenesis in Purkinje cells and motor neurons.

Here, we conducted extensive neuropathological analyses of MATR3 S85C KI mice during early disease stages to determine the temporal emergence of neuropathology and to identify the earliest changes in the properties of the mutant MATR3 protein. We found that MATR3 loss in both Purkinje cells and alpha motor neurons starts to occur during early postnatal stage preceding Purkinje cell degeneration and neuromuscular defects. To understand the immediate downstream consequences of this MATR3 loss, we performed RNA-sequencing (RNA-Seq) in the cerebellum and motor neuron-specific Translating Ribosome Affinity Purification-Sequencing (TRAP-Seq) at early postnatal stages. We identified early yet distinct gene expression changes in the cerebellum and motor neurons, providing insight into the neurodegenerative pathways driven by the pathogenic effects of the S85C mutation in two different motor-controlling neurons, which may inform new cell-specific therapeutic strategies.

## RESULTS

### Purkinje cell loss and neuromuscular junction defects occur after the onset of motor symptoms in MATR3 S85C KI mice

MATR3 S85C KI (*Matr3^S85C/S85C^*) mice develop motor function impairments at 10 weeks of age, which we define as the symptom onset stage (**Figure 1A**)^35^. Their motor function progressively deteriorates, culminating in severe motor impairment by 60 weeks of age (disease end stage), when severe Purkinje cell loss and neuromuscular junction defects are also evident^35^. However, the onset and temporal progression of these neuropathological changes have not been fully delineated. To establish the temporal sequence of pathology, we systematically evaluated Purkinje cell and neuromuscular junction integrity across disease progression.

**Figure 1.**
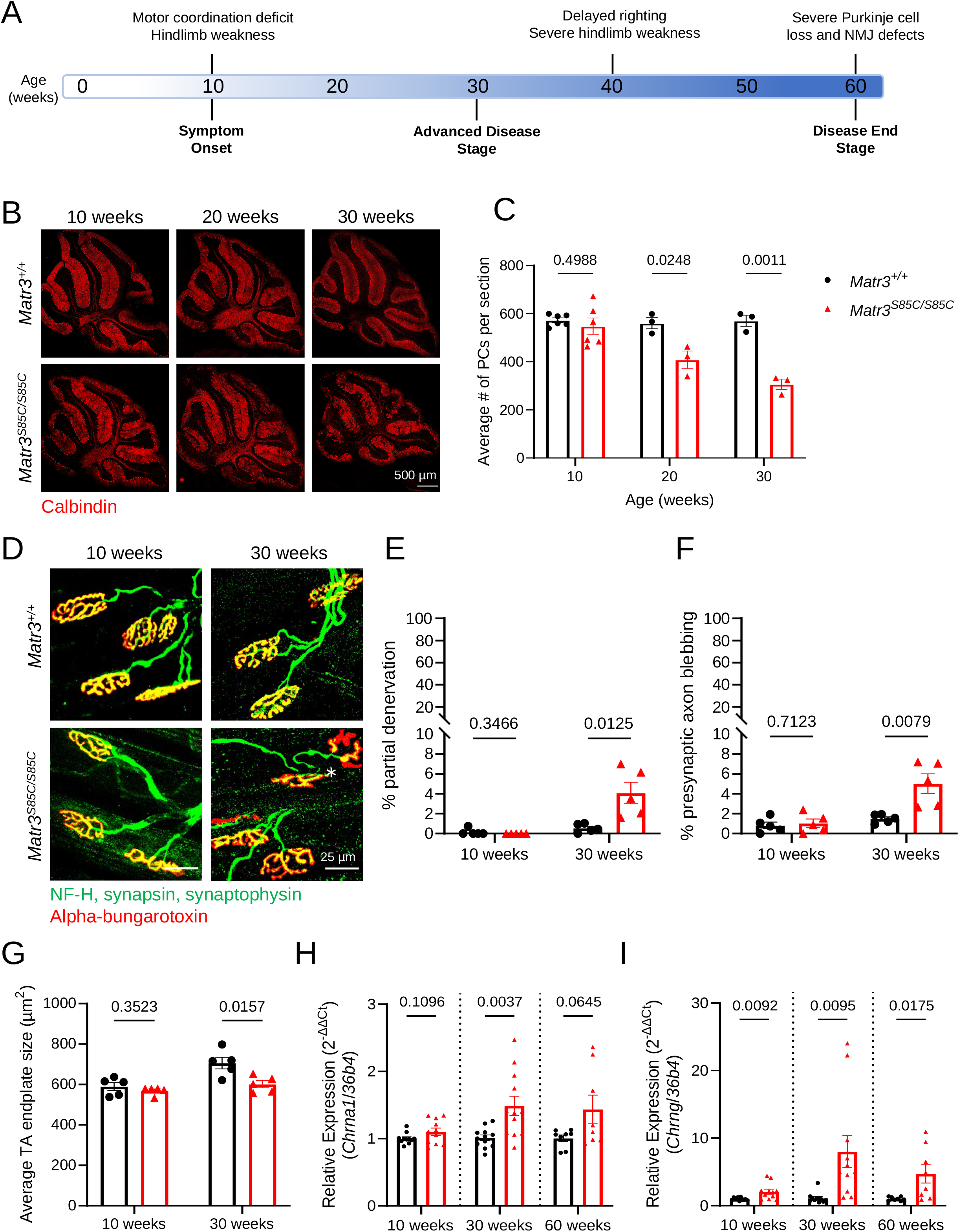
Age-dependent progressive degeneration of Purkinje cells and motor neurons in MATR3 S85C KI mice. **(A)** Timeline of disease progression of MATR3 S85C KI mice (*Matr3^S85C/S85C^*). **(B)** Representative images of Calbindin-positive Purkinje cells in the whole cerebellum of *Matr3^+/+^*and *Matr3^S85C/S85C^* mice at 10, 20, and 30 weeks of age. Scale bars indicate 500 μm. **(C)** Quantification of the number of Purkinje cell soma throughout the whole cerebellum at 10, 20, and 30 weeks of age. Each dot represents the average of 4 sections of a single animal; N=3-6 mice per genotype. **(D)** Representative images of the neuromuscular junctions (NMJs) at 10 and 30 weeks of age. The presynaptic terminal (stained using Neurofilament H, synapsin, and synaptophysin) is shown in green and the motor endplate (α-bungarotoxin-positive staining which targets acetylcholine receptor) in red. Denervated NMJs are indicated by a white asterisk. **(E)** Quantification of partial denervation at the NMJs. Denervation was quantified by partial overlap (>1%, <80%) between the red and green channels. N=5 mice per genotype. **(F)** Quantification of axonal blebbing, which was determined by the presence of thick, rotund staining at the presynaptic terminal. N=5 mice per genotype. **(G)** Quantification of the motor endplate area. A single dot represents the average of >100 NMJs of a single animal; N=5 mice per genotype. **(H-I)** Relative expression (ΔΔC_t_ analysis) of *Chrna1* and *Chrng* from total RNA extracted from tibialis anterior (TA) muscle of *Matr3^+/+^*and *Matr3^S85C/S85C^* mice at 10, 30, and 60 weeks of age as measured by quantitative RT-PCR (N=8-12, male and female mice combined, bar heights depict mean ± SEM with each dot representing a single animal, significance determined by multiple unpaired t-test).

To determine when Purkinje cell loss begins, we conducted cerebellar immunostaining at multiple timepoints from symptom onset (10 weeks of age) to later symptomatic stages (20 and 30 weeks of age). At 10 weeks of age, Purkinje cell numbers were comparable between *Matr3^+/+^*and *Matr3^S85C/S85C^* mice (**Figure 1B, C**). However, by 20 weeks of age, the number of Purkinje cells in *Matr3^S85C/S85C^* mice significantly declined by about 27% compared to the wildtype mice. This degeneration progressed further to an approximately 46% loss by 30 weeks of age (**Figure 1B, C**) and to a ∼70% loss by the disease-end stage^35^. These results indicate that the loss of Purkinje cells is a gradually progressing feature of the disease that occurs after the onset of motor symptoms.

Similarly, to determine when NMJ defects start to develop in the MATR3 S85C KI mice, we first investigated the onset of NMJ pathology in the tibialis anterior (TA) muscle at 10 and 30 weeks of age. At 10 weeks, we did not detect any significant NMJ pathology as evidenced by the lack of denervation and presynaptic axonal blebbing in the NMJs of *Matr3^+/+^* and *Matr3^S85C/S85C^*mice as well as by the comparable size of the TA motor endplate area (**Figure 1D-G**). In contrast, by 30 weeks of age, *Matr3^S85C/S85C^* mice exhibited a modest but significant increase in the percentage of NMJs with partial denervation and axonal blebbing (∼7.4 times and ∼3.4 times that of WT, respectively), along with a statistically significant reduction in TA endplate area when compared to *Matr3^+/+^* mice (**Figure 1D-G, Supplementary Figure 1)**. To complement these morphological analyses, we measured *acetylcholine receptor subunit α1* (*Chrna1*) and *acetylcholine receptor γ-subunit* (*Chrng*) mRNA expression in TA muscles as the upregulation of these genes is a well-established marker of denervation^55-57^. Consistent with the histological findings, *Chrna1* and *Chrng* mRNA levels were markedly elevated in the TA muscles of *Matr3^S85C/S85C^* mice at symptomatic stages (30 weeks and 60 weeks of age), while little to mild elevation of denervation markers (*Chrna* and *Chrng*, respectively) was observed at the symptom onset stage (10 weeks of age) (**Figure 1H, I**).

Collectively, these results demonstrate that both Purkinje cell loss and neuromuscular junction pathology develop progressively and become evident only at later stages of the disease progression after the onset of motor deficits. Thus, we sought to identify earlier neuropathological phenotypes that may contribute to disease progression and potentially represent the initial triggers of pathogenesis.

### Early and progressive loss of MATR3 immunoreactivity precedes the onset of motor deficits and neuropathology

We previously demonstrated that loss of MATR3 immunoreactivity (hereafter, MATR3 loss) is a prominent feature in the majority of Purkinje cells and spinal *α*-motor neurons of the MATR3 S85C KI mice^35, 58^ at advanced disease stages, indicating the selective impact of the mutation in these vulnerable neurons governing motor function. Additionally, we found that extensive loss of MATR3 in Purkinje cells was already evident at 6 weeks of age, prior to motor symptom onset^35^, raising the questions of when this loss starts to occur and whether a similar presymptomatic loss of MATR3 occurs in spinal *α*-motor neurons. Thus, we reasoned that defining the timing of the onset of MATR3 loss in these neurons would help delineate when the mutation first begins to exert its effects and reveal early molecular processes that may potentially underlie neurodegeneration.

To determine the onset and progression of MATR3 loss in these vulnerable motor-associated neuronal populations, we performed immunostaining for MATR3 in combination with cell-type markers (Calbindin for Purkinje cells; ChAT and NeuN for motor neurons) on cerebellar and lumbar spinal cord sections collected from *Matr3^+/+^* and *Matr3^S85C/S85C^* mice at multiple presymptomatic timepoints. At 2 weeks of age, there was no significant difference in the proportion of Purkinje cells with reduced MATR3 staining in both male and female *Matr3^S85C/S85C^* mice compared to the age-matched *Matr3^+/+^* littermates (**Figure 2A-C**). However, by 3 weeks of age, approximately 30-40% of Purkinje cells in *Matr3^S85C/S85C^*mice exhibited a clear reduction in MATR3 staining. This proportion increased further to approximately 60% by 4 weeks of age (**Figure 2A-C**), reached ∼80% at 6 weeks of age^35^, and remained consistently high, exceeding 80%, throughout disease progression (10, 20, and 30 weeks in **Supplementary Figure 2**; 60 weeks in Kao et al., 2020^35^). Notably, Purkinje cell numbers remain unaffected at these early timepoints (**Supplementary Figure 3**), indicating that the early and rapidly progressing depletion of MATR3 precedes detectable neuronal loss and the onset of motor symptoms and may represent an initiating molecular event.

**Figure 2.**
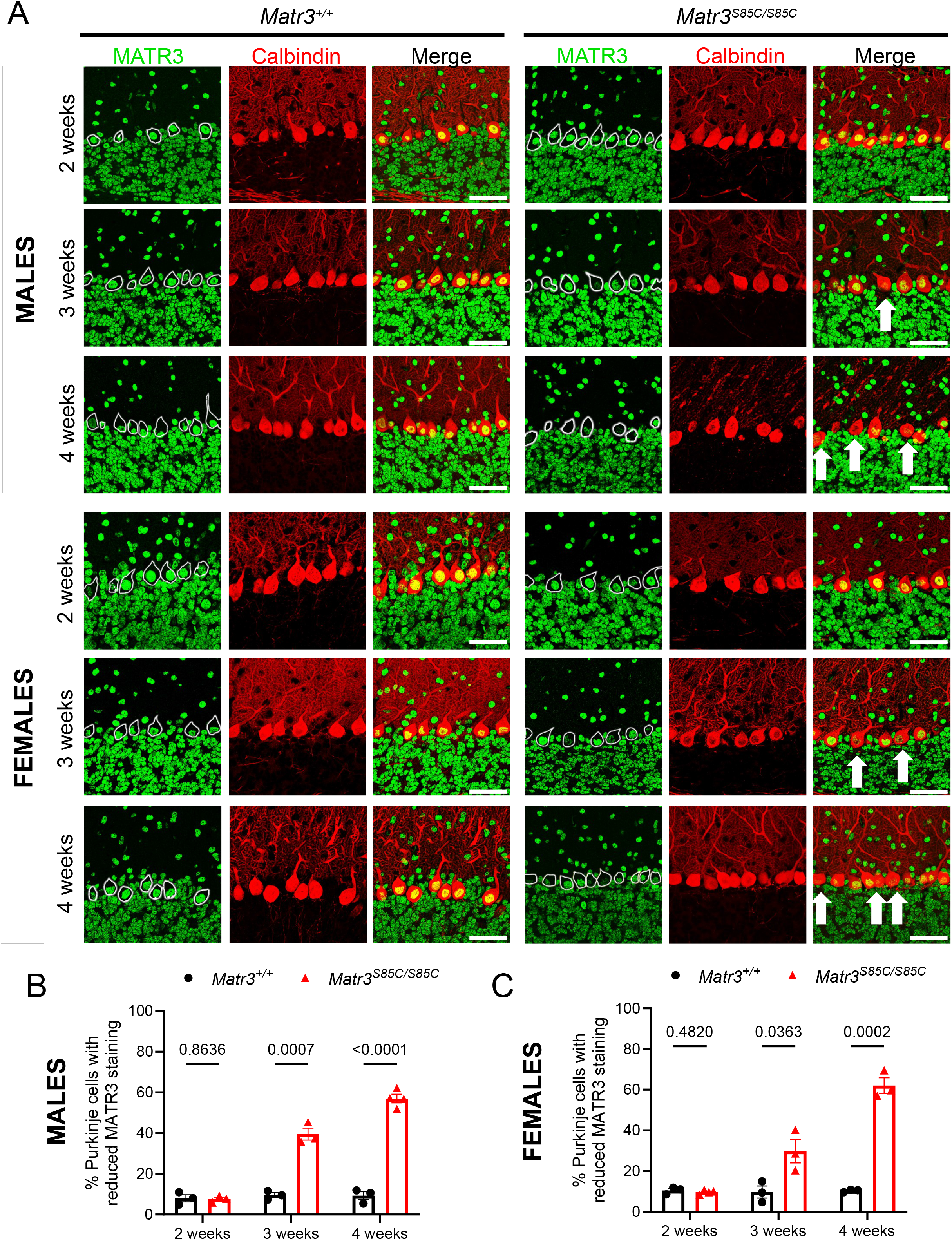
Early and progressive loss of MATR3 in Purkinje cells in MATR3 S85C KI mice. **(A)** Representative images of MATR3 and calbindin staining in the Purkinje cell layer of male and female *Matr3^+/+^* and *Matr3^S85C/S85C^* mice at presymptomatic stages (2, 3 and 4 weeks of age). Examples of Purkinje cells with reduced MATR3 staining are indicated by white arrows. Purkinje cell soma were outlined based on Calbindin immunostaining and are overlayed into the MATR3 channel. **(B, C)** Quantification of the percentage of Purkinje cells throughout the cerebellum with reduced MATR3 staining at 2, 3 and 4 weeks of age in male (**B**) and female (**C**) mice. A single dot represents the average of 4 sections of a single animal; n=3-6 mice per genotype.

In the lumbar spinal cord, there was no significant loss of MATR3 staining in the α- or γ-motor neurons of *Matr3^S85C/S85C^* mice at 4 weeks of age when compared to *Matr3^+/+^* littermates (**Figure 3A-C, Supplementary Figure 4A**). However, by 5 weeks of age, significant MATR3 loss (∼15-30% of cells) became evident specifically in α-motor neurons, the motor neuron subtype that tends to degenerate in ALS, but not in γ-motor neurons, which are spared in ALS^59^. Furthermore, the proportion of α-motor neurons exhibiting reduced MATR3 staining further increased to ∼25-35% at 6 weeks of age while the proportion of γ-motor neurons with reduced MATR3 staining remained comparable between *Matr3^+/+^* and *Matr3^S85C/S85C^* mice (**Figure 3A-C, Supplementary Figure 4A**). This proportion further progressed to ∼70% of α-motor neurons at 10 weeks of age (disease onset) and >80% at 30 weeks of age (advanced disease stage)^58^, demonstrating a progressive disease stage-dependent increase. Importantly, overt loss of motor neuron soma was not observed in *Matr3^S85C/S85C^*spinal cord at these early presymptomatic timepoints **(Supplementary Figure 4B, C)**, supporting the conclusion that MATR3 loss occurs prior to motor neuron phenotypes. Intriguingly, MATR3 loss seems to occur earlier and progressed more rapidly in Purkinje cells compared to α-motor neurons, which is reflected in the pathological manifestations for each cell type, with Purkinje cells degenerating more severely compared to α-motor neuron soma at the disease end stage^35^.

**Figure 3.**
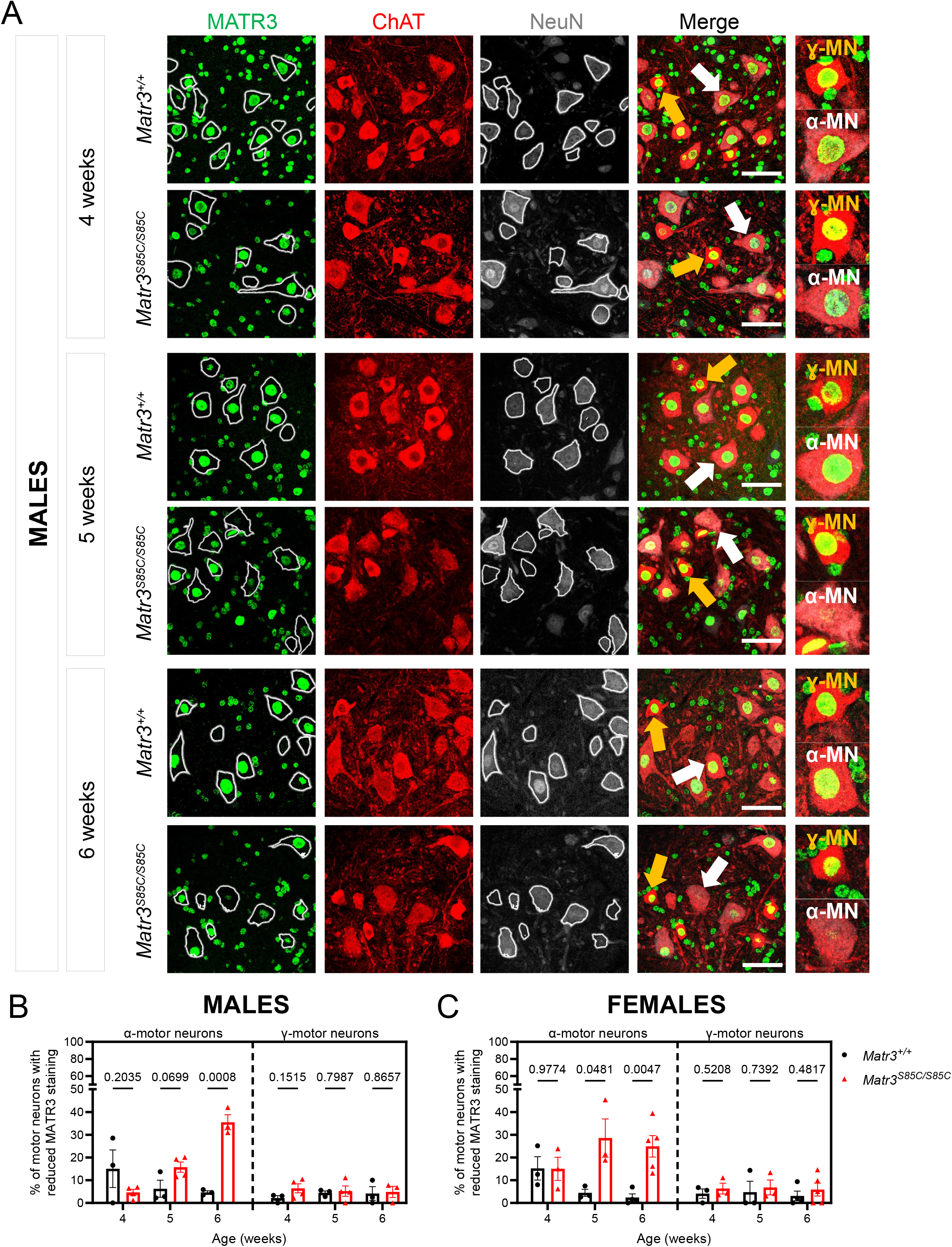
Early and progressive loss of MATR3 in alpha but not gamma motor neurons of MATR3 S85C KI mice. **(A)** Representative images of MATR3 staining in the lumbar ventral horn of male *Matr3^+/+^* and *Matr3^S85C/S85C^* mice, co-stained with ChAT and NeuN. Motor neuron soma were outlined based on ChAT immunostaining and are overlayed into the MATR3 and NeuN channels. α-motor neurons are marked by high staining of ChAT and NeuN, whereas γ-motor neurons are marked by high staining of ChAT and low NeuN. Representative α- and γ-motor neurons are marked by white and yellow arrows, respectively and are shown magnified in the inset. **(B, C)** Quantification of the percentage of α-motor neurons or γ-motor neurons with reduced MATR3 staining at 4, 5, and 6 weeks of age in male (**B**) and female (**C**) mice. A single dot represents the average of 3-4 sections of a single animal; n=6-8 mice per genotype.

Taken together, these findings indicate that MATR3 loss is an early and progressive pathological marker in both Purkinje cells and spinal α-motor neurons in MATR3 S85C KI mice. Its occurrence prior to neurodegeneration and the onset of motor symptoms implicates MATR3 loss as a primary, upstream event that may drive disease initiation.

### Temporal transcriptomic profiling of the cerebellum identifies early dysregulation of Purkinje cell-enriched genes

To investigate the downstream molecular consequences of MATR3 loss in these vulnerable neuronal populations, we next performed transcriptomic profiling to identify early gene expression changes that could represent disease-initiating processes.

First, we focused on characterizing the molecular profile of the cerebellum, where MATR3 loss is specifically found in Purkinje cells, but not in neighboring neurons. Previously, we conducted bulk RNA-sequencing (RNA-seq) on cerebella from *Matr3^S85C/S85C^* and *Matr3^+/+^* mice at the early disease stage (8-10 weeks old) and found 110 differentially expressed genes^35^. To determine the cellular origins of these transcriptomic alterations in our model, we mapped differentially expressed genes identified from this bulk cerebellar RNA-seq dataset to a previously published single-cell RNA-seq atlas of the mouse cerebellar cortex^60^. Consistent with our previous findings, this analysis revealed that genes significantly upregulated in *Matr3^S85C/S85C^* cerebella were predominantly expressed in immune cell populations such as microglia and macrophages **(Figure 4A, B)**. In contrast, genes that were significantly downregulated in *Matr3^S85C/S85C^* cerebella showed preferential expression in Purkinje cells **(Figure 4A, B)**, suggesting that in addition to immune-related cell types, Purkinje cells are among the major affected cell types in the cerebella of these mutant mice.

**Figure 4.**
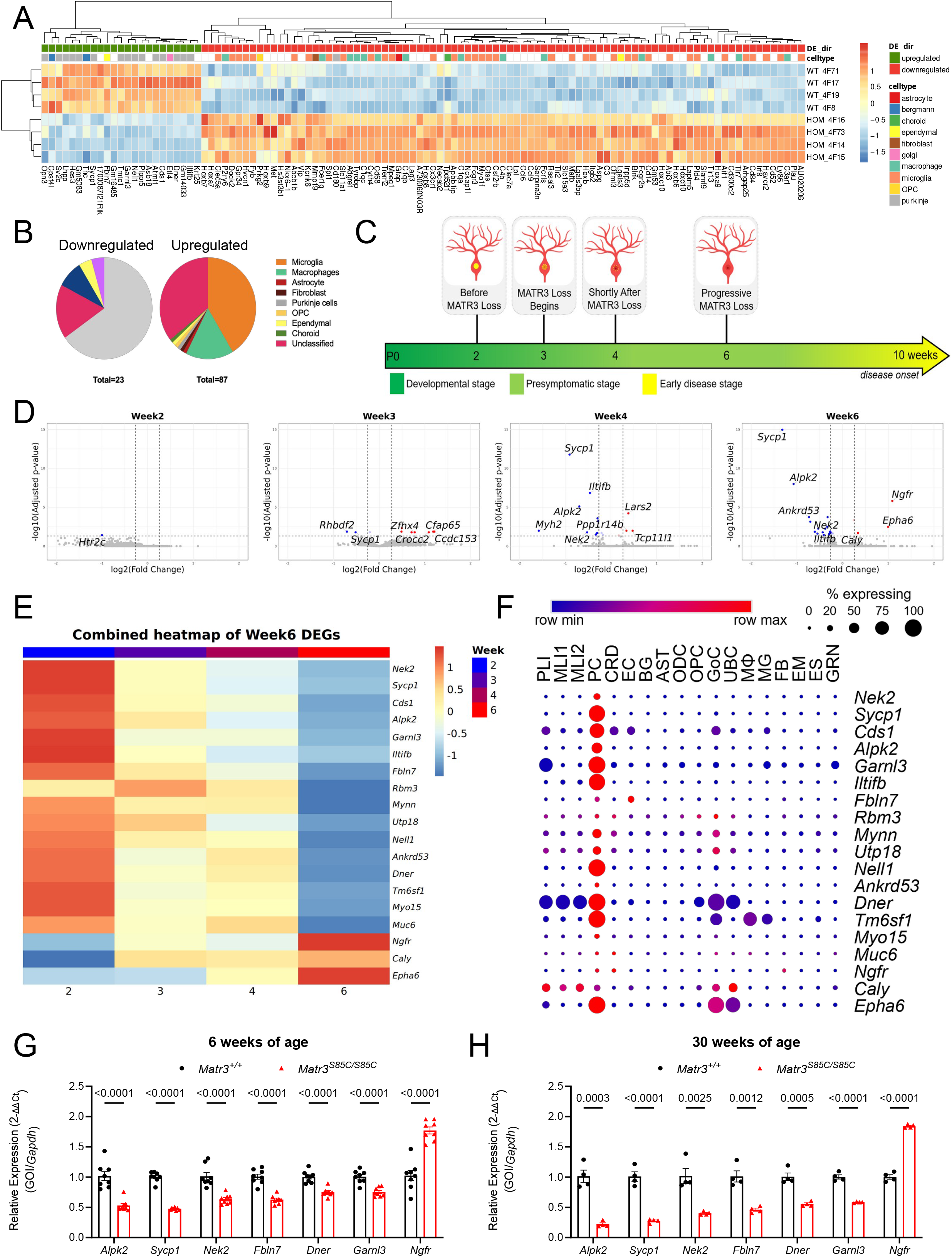
Purkinje cell-enriched gene dysregulation precedes motor symptom onset and neuropathology. **(A-B)** Gene expression heatmap of DEGs identified in mouse cerebellum at 8-10 weeks of age (Kao et al., 2020). Expression values are scaled across samples. Direction of DE is indicated. DEGs that are identified as cell-type specific gene markers in the cerebellum in adult mouse cerebellum are indicated (Kozareva et al., 2021). Pie charts illustrating the proportion of 8-10-week DEGs identified as a marker for specific cell types. **(C)** Timeline showing the timing of bulk cerebellar RNA-seq with regards to the onset of MATR3 loss in the Purkinje cells of MATR3 S85C mice. **(D)** Volcano plots showing differentially expressed genes in the cerebellum of *Matr3^+/+^* and *Matr3^S85C/S85C^* mice at 2, 3, 4, and 6 weeks of age. The number of DEGs increased over time, consistent with progressive MATR3 loss in Purkinje cells. **(E)** Heatmap showing the expression profiles of genes identified as differentially expressed at Week 6 across the Week 2, 3, 4, and 6 cerebellar RNA-seq datasets. Expression values are scaled across samples. **(F)** Cell-type specific expression in the adult mouse cerebellum (Kozavera et al., 2021) of DEGs identified at 6 weeks. **(G-H)** Relative expression (ΔΔC_t_ analysis) of select 6 week-DEGs from total RNA extracted from the cerebellum of *Matr3^+/+^* and *Matr3^S85C/S85C^* mice at 6 and 30 weeks of age as measured by quantitative RT-PCR (N=3-9 mice per genotype, bar heights depict mean ± SEM with each dot representing a single animal, significance determined by multiple unpaired t-test).

To capture the early molecular disturbances preceding overt pathology and potentially contributing to Purkinje cell pathology, we harvested cerebella from *Matr3^S85C/S85C^* and *Matr3^+/+^* mice at 2, 3, 4, and 6 weeks of age and conducted bulk cerebellar RNA-seq **(Figure 4C)**. Given potential sex-specific molecular differences, this analysis was restricted to female mice as was previously performed for 8-10 weeks. Notably, these presymptomatic timepoints encompass the period before and after the initiation of significant MATR3 loss in Purkinje cells, thereby providing a temporal window to identify downstream transcriptional changes associated with this loss, before the emergence of functional deficits. Furthermore, these timepoints capture a progressive gradient of MATR3 loss in Purkinje cells, ranging from minimal loss (<10% in 2 weeks old) to extensive depletion (∼80% in 6 weeks old), enabling the characterization of transcriptional responses across different stages of MATR3 deficiency.

Few differentially expressed genes (DEGs) were identified in the cerebellum in *Matr3^S85C/S85C^* mice at 2, 3, 4 and 6 weeks of age compared to *Matr3^+/+^* littermates (**Figure 4D**). We identified a single DEG at 2 weeks, indicating that molecular perturbations in the cerebella are initially subtle, consistent with the absence of detectable MATR3 loss at this stage. Starting at 3 weeks, when MATR3 loss first becomes detectable, and continuing through later ages, the number of DEGs progressively increased, accompanied by greater magnitudes of fold change with increasing age (**Supplementary Figure 5**). The heatmap in **Figure 4E** shows temporal changes in gene expression of week 6 DEGs as progressive loss of MATR3 occurs. Among them, *Sycp1* (synaptonemal complex protein 1), which is highly enriched in Purkinje cells, was consistently downregulated across the 3-, 4-, and 6-week datasets with the extent of the downregulation increasing with increased MATR3 loss. A few other genes that are downregulated in both 4 and 6 weeks were *Iltifb* (Interleukin-10-related T-cell-derived-inducible factor beta), *Nek2* (NIMA-related kinase 2), and *Alpk2* (Alpha kinase 2). At 6 weeks, when the majority of Purkinje cells exhibit MATR3 loss, we found several strongly upregulated genes including *Ngfr* (Nerve growth factor receptor) and *Epha6* (EPH receptor A6), which have been previously implicated in neuronal stress, cell death, and pathological conditions^61, 62^. Given the emergence of these stress-induced, cell death-associated genes at 6 weeks, we therefore focused on examining the DEGs identified at this time point.

Consistent with our analysis of 8-10 week (symptom onset) cerebella (**Figure 4A, B**), the majority of downregulated DEGs identified at 6 weeks were predominantly expressed in Purkinje cells **(Figure 4F)**. Notably, in contrast to our findings at symptom onset (**Figure 4A, B)**, changes in the expression of neuroinflammation-related genes were minimal at these early timepoints (**Supplementary Figure 6**), reinforcing the notion that Purkinje cells are among the earliest and most affected cell populations in the cerebellum of the mutant mice. This may suggest that early Purkinje cell transcriptional perturbations may represent disease-initiating events rather than secondary consequences of cell death or inflammation.

To validate the RNA-seq findings, we performed RT-qPCR analysis and confirmed the significant upregulation of *Ngfr* together with the downregulation of *Sycp1*, *Alpk2* and other genes identified at 6 weeks of age **(Figure 4G)**. Notably, these transcriptomic perturbations persisted even at 30 weeks of age **(Figure 4H)**. Together, these validation experiments support the reliability of the transcriptomic data and underscore the early molecular alterations that precede Purkinje cell loss.

### NGFR signaling as an early disease-associated pathway in Purkinje cells

Among the early dysregulated genes, we selected *Ngfr* (also known as p75NTR, p75, CD271, LNGFR, TNFRSF16) for further downstream characterization because it was among the most highly upregulated genes **(Figure 4F)** and has been implicated in neuronal cell death pathways^62-64^. NGFR is a disease- and injury-induced gene that is normally expressed during development and reactivated in the adult brain under pathological conditions^65-67^. In wildtype cerebella, among the timepoints examined, we found that *Ngfr* expression was highest at postnatal week 2 and was subsequently and consistently suppressed at postnatal weeks 3, 4, and 6 (**Figure 5A**). In contrast, *Ngfr* expression was significantly induced in mutant cerebella at 6 weeks of age and remained persistently upregulated even at 30 weeks of age (**Figures 4G-H and 5A**), suggesting its re-expression may occur in response to extensive MATR3 loss. Notably, this transcriptomic upregulation is also reflected in the protein levels as we found that NGFR is profoundly upregulated in cerebellar lysates in both male and female mutant mice at 9-11 weeks of age **(Figure 5B-C)**. To determine whether increased NGFR expression was Purkinje cell-specific, we performed immunostaining on cerebellar sections from *Matr3^S85C/S85C^* mice and *Matr3^+/+^* mice with NGFR and Calbindin antibodies. Similar to previous reports^68, 69^, we found that NGFR was restricted to Purkinje cells in specific lobules, VI/VII and IX/X in wildtype mice (**Figure 5D**). In the mutant mice, NGFR immunoreactivity was increased throughout the cerebellum with greater elevation in lobules VI/VII and IX/X (**Figures 5D-E**). To determine whether NGFR signaling is mediated by the activation of the JNK pathway, we performed immunostaining for phospho-c-JUN (p-c-JUN), which is known to localize to the nucleus upon activation^70-72^. Notably, we found increased number of Purkinje cells with nuclear p-c-JUN signals in the cerebellum of the mutant mice at 15 weeks **(Figure 5F-G and Supplementary Figure 7)**, which preceded the onset of significant Purkinje cell loss at 20 weeks of age. In contrast, no nuclear p-c-JUN-positive Purkinje cells were detected at 5 weeks of age **(Figure 5F, G)**. Furthermore, we examined the levels of downstream cell death-related genes activated by the NGFR-JNK pathway. We found that *Bax* and *Caspase 3* showed increased expression in the mutant mice **(Figure 5H).** In parallel, given the context-dependent roles of NGFR in both survival and cell death pathways^73^, we also assessed the expression of NGFR-NFκB-associated cell survival genes. Intriguingly, the expression of pro-survival genes such as *Bcl2, Mcl1,* and *Xiap* were not significantly altered **(Figure 5H)**, supporting the possibility that NGFR signaling in the mutant cerebellum may preferentially engage cell death-associated pathways. Altogether, these findings identify aberrant NGFR signaling as an early disease-associated molecular response that precedes Purkinje cell loss and may contribute to Purkinje cell degeneration.

**Figure 5.**
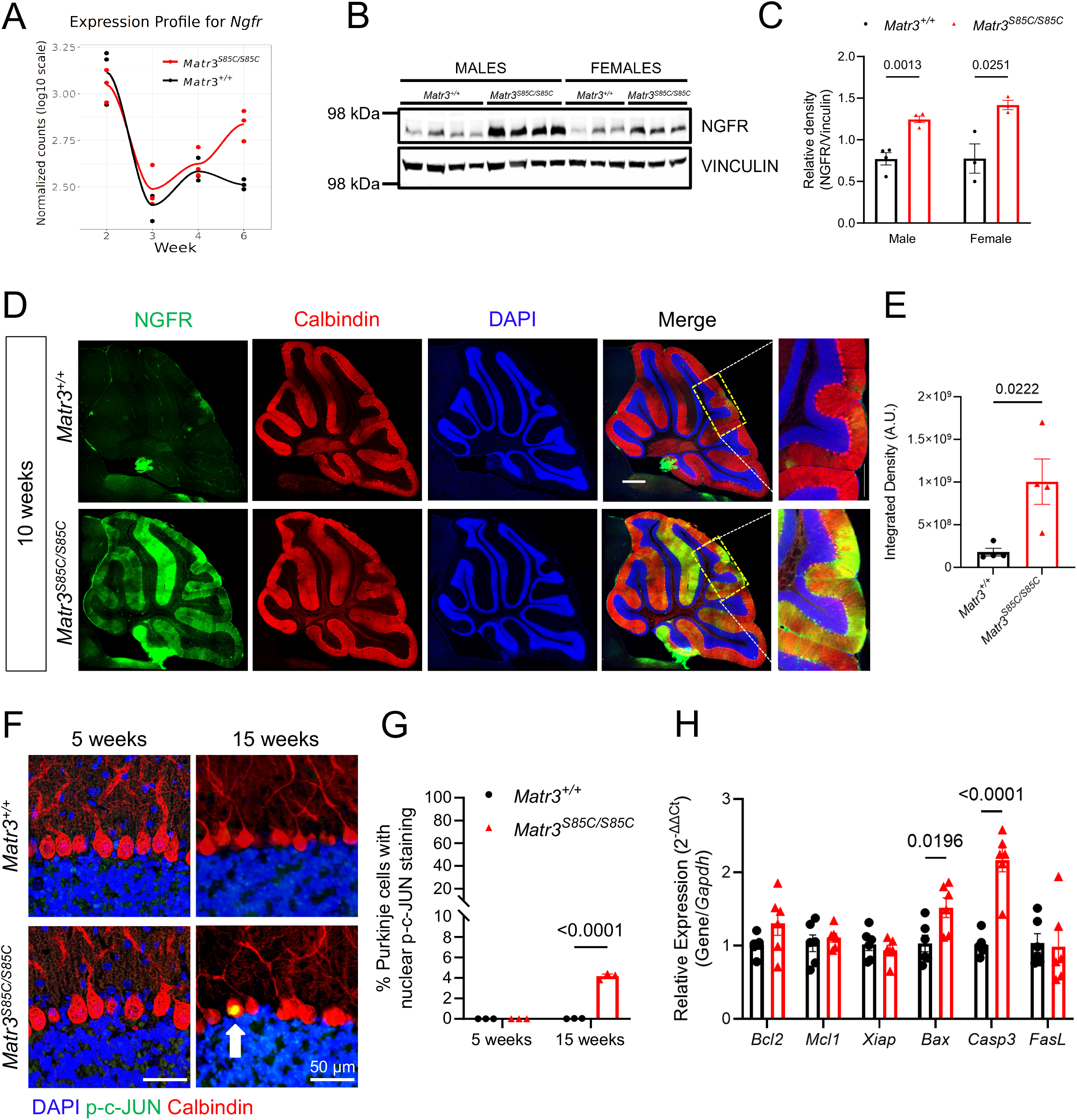
Early activation of the NGFR-JNK signaling pathway in Purkinje cells of MATR3 KI mice. **(A)** Expression of *Ngfr* in the cerebellum of *Matr3^+/+^*and *Matr3^S85C/S85C^* mice at 2, 3, 4, and 6 weeks of age as determined by RNA-seq. **(B, C)** Immunoblot and quantification for NGFR and VINCULIN from protein lysates obtained from the cerebellum of 9-week-old *Matr3^+/+^* and *Matr3^S85C/S85C^*mice. **(D)** Representative images of the cerebellum of MATR3 S85C KI mice and wildtype littermates immunostained for NGFR (green), Calbindin (red), and DAPI (blue) at 10 weeks of age. **(E)** NGFR intensity is significantly increased in the cerebellum of MATR3 S85C KI mice at 10 weeks of age. Each dot represents average integrated density of 4 sections of one animal; n=4 mice per genotype. **(F)** Representative images of calbindin-positive Purkinje cells with nuclear phospho-c-JUN (p-c-JUN) staining at 5 and 15 weeks of age. Purkinje cells with nuclear p-c-JUN staining are indicated by a white arrow. **(G)** Quantification of the percentage of Purkinje cells with nuclear p-c-JUN. A single dot represents the average of 4 sections of a single animal; n=3 mice per genotype. **(H)** Relative expression (ΔΔC_t_ analysis) of pro-survival genes *Bcl2, Mcl1, Xiap* and pro-apoptotic genes *Bax, Casp3, Fasl* from total RNA extracted from the cerebellum of *Matr3^+/+^* and *Matr3^S85C/S85C^* mice at 12 weeks of age as measured by quantitative RT-PCR (N=6 mice per genotype, bar heights depict mean ± SEM with each dot representing a single animal, significance determined by multiple unpaired t-test).

### Motor neuron-specific RNA profiling reveals early and progressive transcriptional alterations across disease stages

To identify early and progressive molecular mechanisms underlying motor neuron degeneration in *Matr3^S85C/S85C^*mice, we conducted RNA profiling using the Translating Ribosome Affinity Purification (TRAP) system. This approach enables isolation of mRNA actively engaged with GFP-tagged ribosomes, thereby capturing the translatome of defined cell populations^74, 75^. To selectively profile motor neurons, we utilized transgenic mice expressing EGFP-tagged ribosomal subunit Rpl10a under the control of the *Chat* promoter (*Chat-EGFP/Rpl10a*). We then crossed these TRAP transgenic mice to *Matr3^S85C/+^* mice to obtain *Chat-EGFP/Rpl10a; Matr3^S85C/S85C^* and *Chat-EGFP/Rpl10a; Matr3^+/+^* mice as a control (**Figure 6A**). Successful enrichment of motor neuron transcripts in immunoprecipitated (IP) samples was confirmed by qRT-PCR (**Figure 6B**). Canonical motor neuron markers *Chat* and *Mnx1* were strongly enriched in IP fractions and subsequently de-enriched in the flow through (FT) fractions, whereas astrocytic and microglial markers (*Aldh1l1* and *Cx3cr1*) were markedly de-enriched in the IP fractions, confirming the specificity of the TRAP isolation for motor neurons.

**Figure 6.**
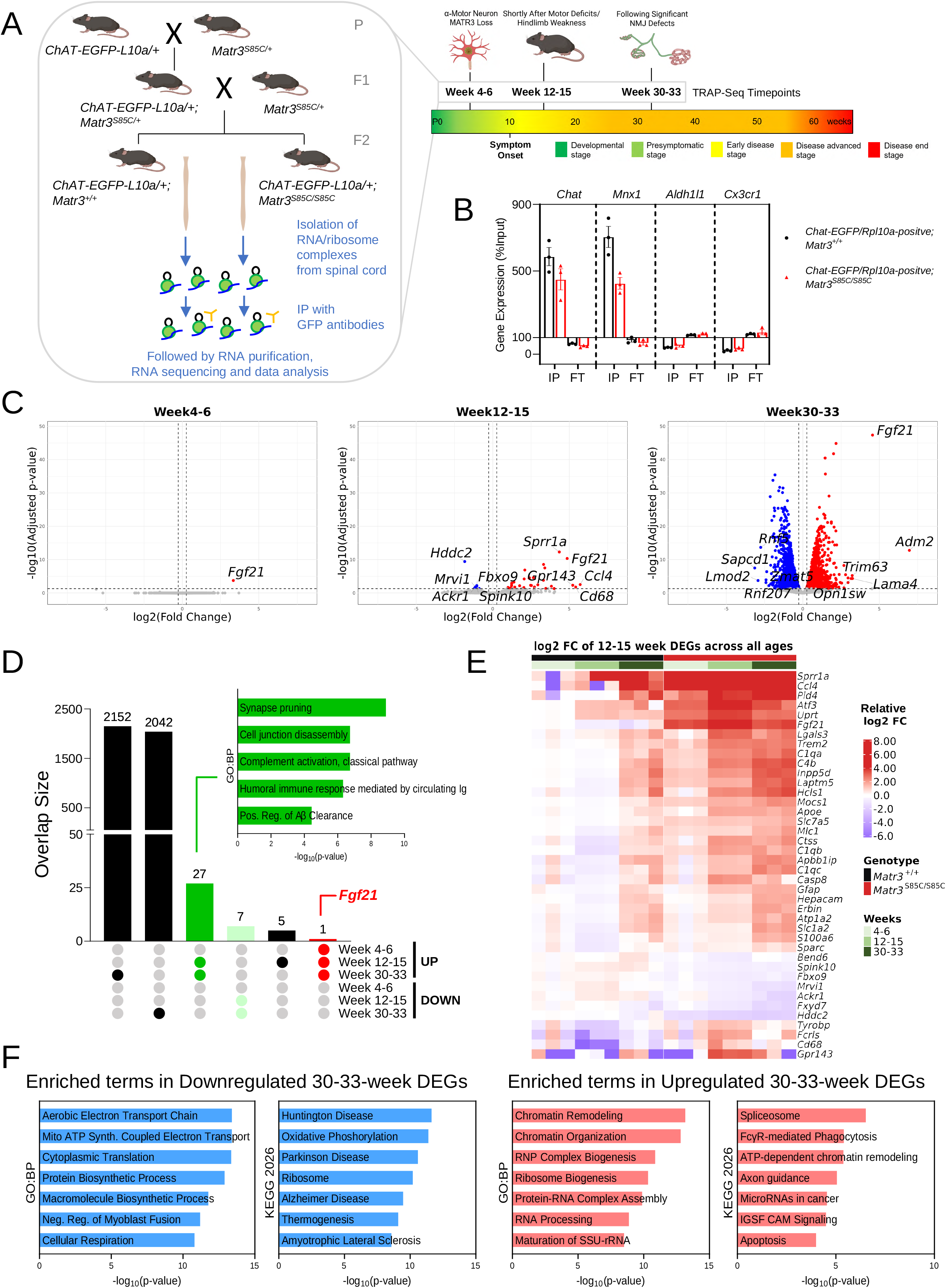
Motor neuron translatome dysregulation emerge and intensify with disease progression. **(A)** Genetic scheme to generate *Matr3^+/+^*; Chat-EGFP/Rpl10a-positive and *Matr3^S85C/S85C^*; Chat-EGFP/Rpl10a-positive mice. RNA from EGFP-positive ribosomes from the spinal cord of each genotype were isolated and sequenced (TRAP-Seq). Timeline of disease progression with TRAP-Seq timepoints and neuropathological and behavioral phenotypes annotated. **(B)** qRT-PCR confirmation of enrichment of motor neuron, astrocytic, and microglial markers in the RNA isolated from the immunoprecipitated (IP) and flowthrough (FT) fractions relative to levels in the input fraction. **(C)** Volcano plot showing differentially expressed genes in the motor neuron translatome of *Matr3^+/+^* and *Matr3^S85C/S85C^* mice at pre-symptomatic (4-6 weeks old), early symptomatic (12-15 weeks old), and at advanced disease (30-33 weeks old) stages. Identified DEGs increases with disease progression. **(D)** UpSetR plots identifying shared and unique DEGs across the disease progression. Gene sets that are consistently upregulated from all three timepoints are shown in red while gene sets that were either consistently upregulated or consistently downregulated during both symptomatic timepoints are shown in green or light green. Inset show gene ontology (biological processes; GO:BP) analysis of the indicated gene set, analysis was performed using Enrichr. **(E)** Heatmaps illustrating gradual dysregulation of early symptomatic DEGs starting from the presymptomatic stage and progressing through the advanced disease stage. Baseline mean expression was calculated from wild-type samples at Weeks 4 to 6, and individual counts across all time points were normalized to this baseline to derive relative log_2_fold changes. **(F)** Pathway enrichment analysis (KEGG, GO:BP) of upregulated (in red) or downregulated DEGs (in blue) identified at the advanced disease stage (30-33 weeks old). Analysis was performed using Enrichr.

Next, we performed TRAP-seq at three key disease stages (**Figure 6A**): a presymptomatic stage which coincides with the onset of MATR3 loss in alpha motor neurons (4-6 weeks of age), early symptomatic stage when motor function begins to decline (12-15 weeks of age), and advanced disease stage immediately following significant NMJ denervation (30-33 weeks of age). This design enabled temporal resolution of motor neuron-specific transcriptomic changes during disease progression. Differential expression analysis (DESeq2; padj < 0.05, log_2_fold change ≥ 0.263 or ≤ -0.263) revealed progressively increased transcriptional perturbations as disease advanced (**Figure 6C**). At the earliest presymptomatic stage (4–6 weeks), *Fgf21* (fibroblast growth factor 21) was the only significantly upregulated gene. The number of differentially expressed genes increased at the early symptomatic stage (12-15 weeks) and this number rose substantially further by the advanced stage (30-33 weeks), indicating progressive disruption of motor neuron gene expression programs with disease progression.

Intriguingly, we found that most genes that are dysregulated at the early symptomatic stage (12– 15 weeks, 28/33 upregulated genes and 7/7 downregulated genes) remained persistently perturbed in the same direction at the advanced disease stage (30-33 weeks) (**Figure 6D**). Pathway enrichment analysis of the genes that were co-upregulated at both symptomatic stages revealed an enrichment for biological processes including synapse pruning, cell junction disassembly, complement activation, and several immune response-related terms, suggesting a progressive engagement of immune-related pathways and synaptic remodeling programs within motor neurons during the early stages of disease progression. Further analysis of the DEGs at this early symptomatic stage revealed that the expression of the upregulated DEGs showed trending increase during the presymptomatic stage (4–6 weeks) and remained elevated through the advanced stage (30–33 weeks) (**Figure 6E**). Conversely, genes that are downregulated during disease remained consistently downregulated across subsequent stages. Overall, these results reveal a progressive and sustained perturbation of the motor neuron translatome in the mutant spinal cord.

Given the large number of DEGs at advanced disease stage (30-33 weeks) when NMJ structural defects occur, we performed gene ontology analysis to further define the functional trajectory of these transcriptional changes (**Figure 6F**). Downregulated genes at the advanced disease stage were strongly enriched for pathways associated with mitochondrial function, cellular energy production and protein synthesis. In contrast, upregulated genes at the advanced disease stage were enriched for pathways reflecting immune activation, proteotoxic stress, and apoptosis.

### Early and sustained induction of *Fgf21* and integrated stress response genes precedes overt motor neuron pathology

Among the transcriptional alterations identified by TRAP-seq, the upregulation of *Fgf21* emerged as the earliest and most persistent change during disease progression (**Figures 6C-D and 7A**). *Fgf21* is induced and secreted in response to various acute and chronic cellular stresses and is often upregulated alongside other integrated stress response (ISR) genes following activation of the ISR^76, 77^. Given striking emergence of *Fgf21* expression early in disease, we sought to validate the increase in *Fgf21* levels and investigate whether other ISR genes are similarly upregulated in the motor neurons at presymptomatic stage. qRT-PCR analysis of TRAP-isolated RNA from pre-symptomatic 9-week-old mice revealed robust upregulation of multiple ISR-associated genes, including *Fgf21*, *Gdf15*, *Mthfd2*, *Atf3*, and *Ddit3* (**Figure 7B**). Consistent with this, similar induction of ISR-associated genes was detected in bulk spinal cord mRNA as early as 6 weeks of age (**Figure 7C**) and beyond (9 weeks of age in **Supplementary Figure 8**, 30 weeks of age in **Supplementary Figure 9**). The continuous induction of these ISR-associated genes in combination with the downregulation of genes in protein biosynthetic pathways at the advanced disease stage (**Figure 6F**) suggests that ISR induction represents an early and robust feature of disease in the spinal cord. Notably, at the protein level, ELISA analysis of spinal cord lysates revealed a significant increase in FGF21 abundance at the early disease stage (10-12 weeks) (**Figure 7D**), demonstrating that transcriptional induction of *Fgf21* translates into elevated protein production during disease progression.

**Figure 7.**
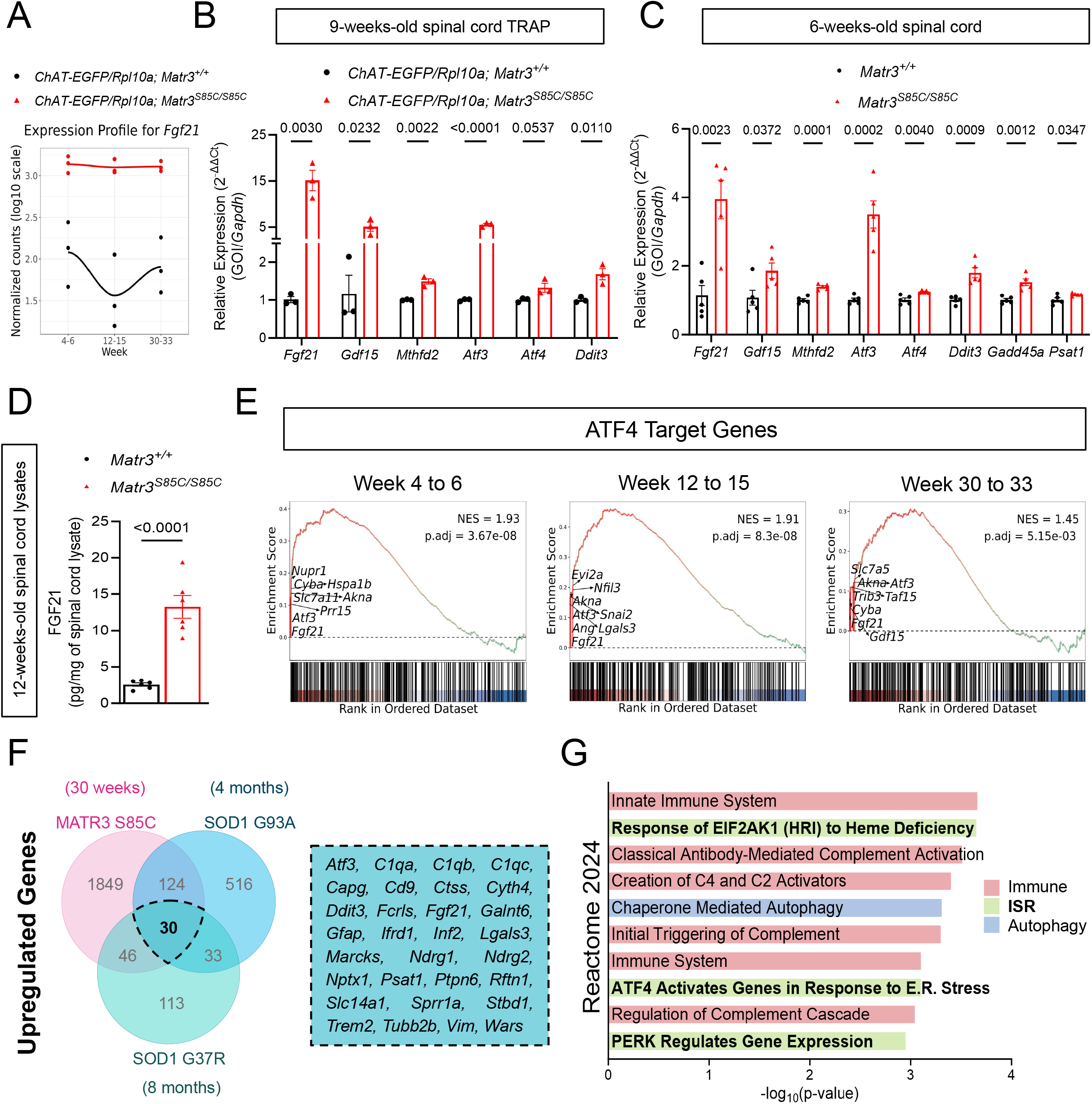
Early and persistent activation of the integrated stress response in the motor neurons of MATR3 S85C KI mice. **(A)** Expression of *Fgf21* across the disease progression as identified by RNA-Seq. **(B, C)** Relative expression (ΔΔC_t_ analysis) of ISR-associated genes from TRAP isolated RNA from the spinal cord of *Chat-EGFP/Rpl10a-positive;Matr3^+/+^* and *Chat-EGFP/Rpl10a-positive;Matr3^S85C/S85C^* mice at 9 weeks of age or total RNA extracted from the spinal cord of *Matr3^+/+^* and *Matr3^S85C/S85C^* mice at 6 weeks of age as measured by quantitative RT-PCR (N=3-5 mice per genotype, bar heights depict mean ± SEM with each dot representing a single animal, significance determined by multiple unpaired t-test). **(D)** FGF21 protein levels in the spinal cord of *Matr3^+/+^* and *Matr3^S85C/S85C^* mice at 12 weeks of age as measured by ELISA. **(E)** Gene set enrichment analysis shows early and persistent positive enrichment of ATF4 target genes in *Chat-EGFP/Rpl10a-positive;Matr3^S85C/S85C^* mice across all disease stages. **(F)** Venn diagram showing overlapping DEGs among upregulated genes identified in ChAT-bacTRAP MATR3 S85C mice at 30-33 weeks of age (this study), ChAT-bacTRAP in SOD1 G93A mice at 4 months of age (Shadrach et al., 2021), and ChAT-RiboTag in SOD1 G37R mice at 8 months of age (Sun et al., 2015). Numbers indicate the total and shared upregulated genes identified in each dataset. Shared upregulated genes are identified in the box. ISR-related genes are colored in red. **(G)** Enrichment analysis (Reactome 2024) of the shared upregulated genes in (F). Analysis was performed using Enrichr. Terms associated with the ISR pathway are bolded and colored in green.

**Figure 8.**
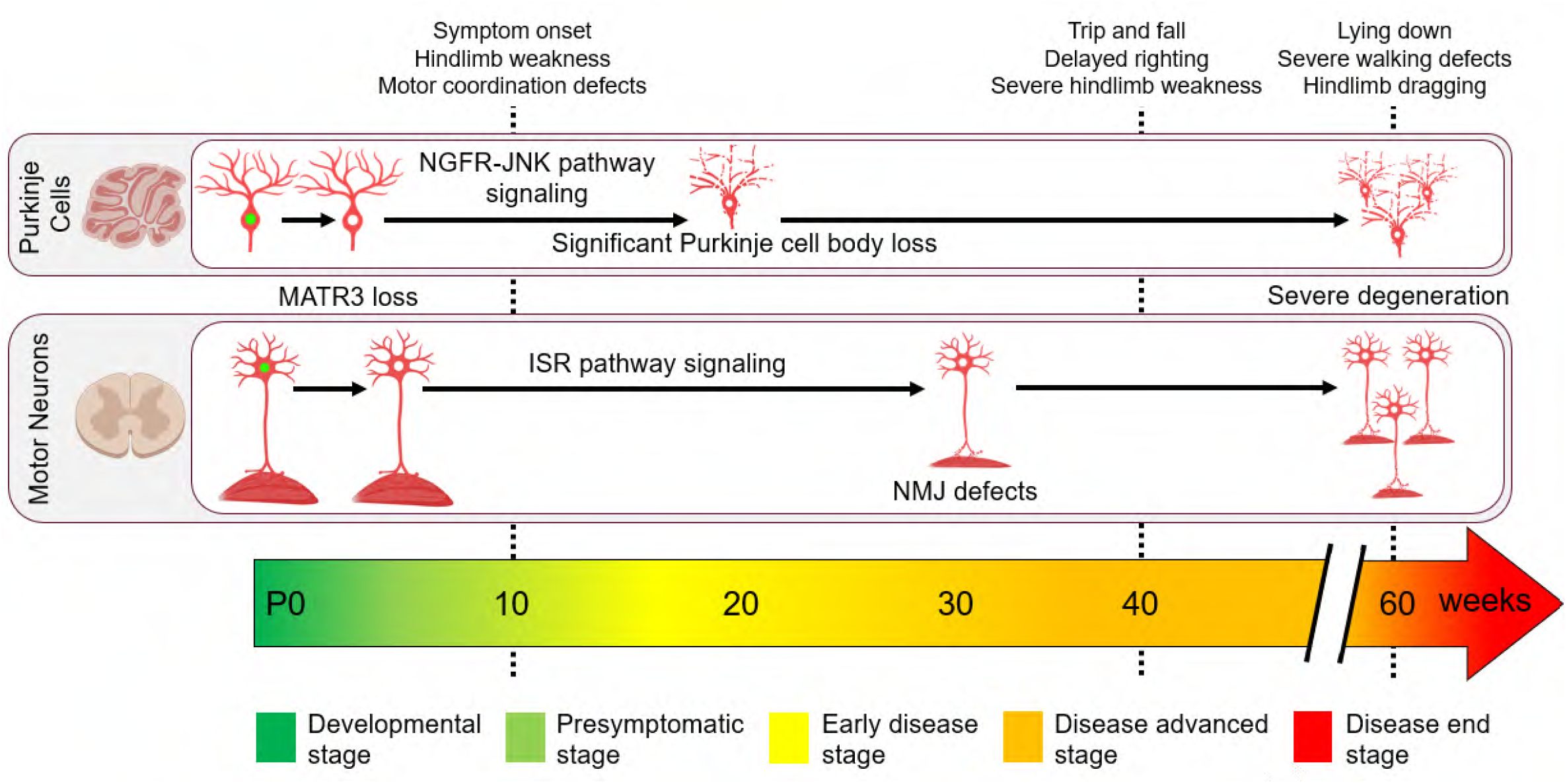
Timeline of neuropathological and motor deficit progression in MATR3 S85C KI mice. The MATR3 S85C mutation selectively affects Purkinje cells and alpha motor neurons, with disease pathogenesis potentially initiated by early postnatal loss of MATR3. Overt neuropathology becomes evident only after disease onset and progressively worsens in parallel with the development of severe motor deficits.

To further validate the induction of the ISR pathway in the motor neurons of the mutant mice, we examined the transcriptomic activity of ATF4, the most characterized downstream effector of ISR, using gene set enrichment analysis (GSEA) against a curated set of ATF4 target genes across all three disease time points. ATF4 target genes were significantly enriched in the mutant motor neurons at all three time points, providing additional evidence of sustained ISR activation throughout disease progression (**Figure 7E**).

To determine whether ISR-associated gene induction represents a shared feature of ALS-related motor neuron stress, we compared the upregulated DEGs we have identified from our dataset to the upregulated DEGs identified from previously published motor neuron-specific transcriptomic datasets of SOD1 mouse models, G93A (Shadrach et al., 2021)^13^ and G37R (Sun et al., 2015)^12^ (**Figure 7F**). This cross-model analysis revealed an overlap of 30 genes that were consistently upregulated across all models, including ISR-upregulated genes *Fgf21*, *Atf3*, *Ddit3, Wars,* and *Psat1*. Gene ontology analysis of these shared upregulated genes reveal an enrichment in ISR- and immune-related pathways (**Figure 7G**). These results suggest that the shared presence of ISR-associated genes in these familial ALS mouse models may reflect a conserved molecular response to disease and supports a potential role for ISR in the initiation and progression of motor neuron degeneration.

### Distinct downstream transcriptomic changes in cerebellum and motor neurons despite shared MATR3 pathology

To determine whether cerebellar neurons and motor neurons share common molecular mechanisms underlying neurodegeneration, we performed a cross-dataset comparison between the 6 week-cerebellar transcriptomic dataset and motor neuron TRAP-seq datasets collected at 30-33 weeks. UpsetR analysis revealed few genes that were commonly dysregulated between the cerebellum and motor neurons in either comparison (**Supplementary Figure 10A**). Since we only identified few DEGs in the cerebellum at 6 weeks of age, we further analyzed coordinated gene expression changes using GSEA. In contrast to the findings in the motor neurons at 4-6 weeks of age, we found no significant enrichment of ATF4 target gene signatures in the cerebellum of 6 weeks-old mutant mice (**Supplementary Figure 10B**). Furthermore, qRT-PCR analysis of ISR-associated genes in the 6-week-old cerebellum failed to recapitulate the upregulation observed in the spinal cord at the same age. Similarly, we found that the dysregulated genes identified from cerebellar transcriptomic dataset such as *Ngfr*, *Casp3,* and *Bax* were not altered in the motor neurons (**Supplementary Figure 10C, D)**. Overall, these results support the notion that the cerebellar/Purkinje cells and motor neuron populations undergo distinct molecular changes during neurodegeneration despite shared MATR3 pathology.

## DISCUSSION

Emerging neuropathological studies of postmortem tissue from ALS patients and animal models are reshaping the understanding of ALS pathology – the impact is not limited to solely the motor neurons but extends to other motor-controlling neurons such as Purkinje cells in the cerebellum. However, little is known regarding the temporal progression of neurodegeneration in motor neurons and Purkinje cells. Particularly, the initial degenerative events and mechanisms remain poorly understood. Furthermore, it remains unclear whether these different types of neurons undergo degeneration through shared or cell type-specific molecular mechanisms.

Our findings reveal that the loss of Purkinje cell soma in MATR3 S85C KI mice is age-dependent, emerging after symptom onset and becoming statistically significant at 20 weeks of age. In addition, NMJ denervation associated with axonal swelling appear long after symptom onset at 30 weeks, but loss of motor neuron soma was not observed until later disease stage (60 weeks)^35^. This suggests that Purkinje cells and motor neurons display different temporal trajectories of neuropathology, with Purkinje cells undergoing relatively early and aggressive pathological changes, while motor neurons exhibit a more gradual course of degeneration. Our findings also reveal that Purkinje cell- and α-motor neuron-specific loss of MATR3 precedes the onset of motor function defects and neuropathology, implicating early postnatal MATR3 loss as an indicator of disease initiation that may potentially trigger the molecular cascade leading to neurodegeneration. Notably, MATR3 loss in Purkinje cells was initiated earlier and progressed to a greater extent than in α-motor neurons, potentially contributing to the earlier onset and greater severity of neuropathological changes in Purkinje cells. Intriguingly, recent proteomic studies have reported significantly reduced levels (∼50%) of MATR3 in motor neurons from sporadic ALS patients^78, 79^, supporting the notion that MATR3 depletion may contribute to motor neuron degeneration outside of MATR3-associated ALS cases.

These collective pathological findings led us to establish the timeline of neuropathological events, enabling us to define a clear window for identifying early molecular changes in Purkinje cells and motor neurons. RNA profiling of the cerebellum and motor neurons at early postnatal stages, prior to the onset of MATR3 loss, revealed few dysregulated genes, further supporting that MATR3 loss may represent an initiating event in disease progression. In contrast, once MATR3 loss began, the number of dysregulated genes increased substantially and increased even further with disease progression. However, it is not yet clear whether these transcriptional dysregulations arise from the disruption of MATR3’s DNA-associated or RNA-associated functions or both. Intriguingly, transcriptomic analyses of the cerebellum and motor neurons identified distinct molecular pathways that may underlie their divergent disease trajectories. How this divergence in the transcriptional responses arise despite shared MATR3 loss remains unclear.

In the cerebellum, transcriptome profiling at the presymptomatic stage revealed that the majority of the differentially expressed genes were highly specific and/or highly enriched in Purkinje cells. In contrast, inflammatory and glial-related genes showed little to no differential expression at this stage and became markedly upregulated only at disease onset^35, 80^. The near absence of inflammatory and glia-related transcriptional changes before disease onset suggests that early pathogenic events are primarily neuronal, whereas neuroinflammatory responses emerge later, coinciding with disease onset. This notion is consistent with earlier studies in SOD1 mouse models, where transcriptomic and genetic analyses similarly suggested that neuronal dysfunction precedes the emergence of neuroinflammatory responses^12, 81^.

Notably, our findings suggest that NGFR is potentially involved in mediating Purkinje cell death in the mutant mice. Specifically, we found significant upregulation of NGFR in Purkinje cells throughout the entire cerebellar lobules. This global increase in NGFR immunoreactivity may reflect a widespread neuronal stress response, potentially involving activation of the JNK-cell death pathway, which could contribute to Purkinje cell death. However, the significance of increased NGFR-JNK-cell death pathway should be further investigated through genetic or pharmacological studies. Previous studies reported increased NGFR expression in spinal motor neurons in postmortem tissue from ALS patients and in SOD1 G93A mouse model at symptomatic stage, when significant motor neuron loss is evident, but not during earlier stages of disease, implicating NGFR as a death-signaling receptor^82^. Similarly, our data revealed that NGFR induction does not occur early in the motor neurons, suggesting that NGFR may not represent an early molecular response in the motor neurons of the mutant mice.

It is possible that other dysregulated genes such as *Sycp1*, *Alpk2*, *Nek2* and *Fbln7*, may contribute to the initiation and progression of Purkinje cell degeneration. SYCP1 (Synaptonemal complex protein 1) is a protein scaffold that forms between homologous chromosomes during meiosis^83, 84^. Although SYCP1 has been extensively studied in spermatocyte development, its role in the central nervous system, particularly in Purkinje cells, has not been investigated. ALPK2 has been implicated in cardiac development and cancer^85, 86^; NEK2 is a cell cycle protein and is linked to cancer^87^; FBLN7 (Fibulin-7) is an extracellular matrix protein^88^; however, their physiological roles in the brain remain largely unexplored. Therefore, further investigation is needed to determine whether their downregulation is an adaptive response or contributes to the onset or progression of Purkinje cell degeneration.

In the spinal cord, our findings reveal that the upregulation of *Fgf21* and other integrated stress response (ISR) genes is one of the earliest molecular responses to MATR3 loss in motor neurons. Our identification of *Fgf21* and a broader ISR-associated transcriptional signature in motor neurons from MATR3 S85C, SOD1 G93A, and SOD1 G37R mice reveals a conserved stress-adaptive program shared across genetically distinct ALS models. Recent studies have reinforced this concept by demonstrating that PGAM5-driven mitochondrial ISR activation constitutes a convergent pathogenic mechanism across multiple ALS subtypes^89^, with the suppression of this pathway delaying disease progression, supporting the idea that ISR signaling may be an active driver of disease. In addition, single-cell transcriptomic studies have identified an early disease-associated motor neuron state that is characterized by activation of stress-response pathways^90^, suggesting that maladaptive stress signaling may represent an early transitional state in vulnerable motor neurons preceding overt neurodegeneration. Consistent with these, ISR activation has also been observed in C9ORF72 models and patient-derived tissues, TDP-43 proteinopathy models, and VAPB-associated ALS^89, 91-93^. Collectively, these findings support a model in which diverse ALS genetic models converge on a shared neuronal stress state characterized by activation of the ISR. However, the functional significance of this increased ISR response, whether early increased ISR expression is protective whereas chronic or sustained ISR expression becomes maladaptive and contributes to neurodegeneration^94, 95^, should be investigated.

Notably, the MATR3 S85C KI model recapitulates this shared ALS molecular signature while maintaining endogenous mutant MATR3 expression and exhibiting a slower, progressive disease course compared with the widely used SOD1 G93A overexpression model. These features provide an extended window to capture early pathogenic mechanisms preceding irreversible neuronal loss, establishing MATR3 S85C KI mice as a physiologically relevant model for studying convergent pathways driving motor neuron degeneration across ALS subtypes and for evaluating therapeutic strategies targeting these shared vulnerability pathways.

We acknowledge several limitations of the present study. Both cerebellar and motor neuron transcriptome profiling were performed using only female samples. Although Purkinje cell-specific and motor neuron-specific MATR3 loss was observed in both males and females at a similar time point and no further distinguishable sex differences were observed in validation experiments, future transcriptome profiling should involve both male and female mice to identify any subtle sex-specific differences. Instead of cerebellar bulk RNA sequencing, Purkinje cell-specific RNA profiling would have been more informative for identifying cell-specific molecular changes, which would have enabled a more direct comparison with the motor neuron transcriptomic dataset. TRAP system using *Chat* promoter does not distinguish between the translatome of alpha vs gamma motor neurons (and also of motor neurons vs cholinergic interneurons). It would be informative to perform motor neuron subtype-specific analyses given that alpha motor neurons are more susceptible to ALS-associated insults compared to gamma motor neurons. It is uncertain whether the early dysregulated genes are direct targets or a response to the cellular stress induced by MATR3 loss. Functional studies targeting NGFR in Purkinje cells or FGF21 or other ISR genes in motor neurons will be necessary to determine the functional significance of these molecular changes in disease pathogenesis.

In conclusion, our study characterized the neurodegenerative course of MATR3 S85C KI mice and identified the onset of MATR3 pathology and neurodegeneration, thereby contributing to the understanding of how MATR3 S85C causes neurodegeneration. Our findings identify MATR3 loss as the earliest detectable pathological event that may contribute to disease initiation. We found that this MATR3 loss is accompanied by distinctive transcriptomic changes in Purkinje cells and motor neurons which may underlie different trajectories of neuropathology. Collectively, our findings provide insights into the initiation of molecular level neurodegenerative process in two different ALS-relevant motor controlling neuronal populations. Thus, the early distinct molecular changes may help identify cell-specific therapeutic targets and inform the development of interventions aimed at slowing or preventing disease progression before irreversible neurodegeneration occurs in ALS.

## Acknowledgements

We thank the members of the Park laboratory and Danilo Shevkoplyas, Young Zhou, and Julie Ruston from Dr. Monica Justice laboratory for insightful discussions on this project. We would like to acknowledge Rintaro Masuda and Alanna Love for their contribution to staining and imaging of cerebellar sections. We appreciate the support from The Centre for Phenogenomics (TCP) for mouse care and generation of histology blocks. We thank the lab of Dr. Ronald Cohn and Developmental & Stem Cell Biology Program for the use of their cryostat and SickKids imaging facility for the use of microscopes. This study was supported by Canadian Institutes of Health Research (CIHR, 202104PJT-462444-NSB-CEAB-275899), the Canada Research Chairs Program (CRC-2021-00063) to J.P. Supported in part by USDA/ARS grant CRIS 3092-51000-065-003S and Research Vision at Texas Children’s Hospital to H.K.Y. K.M. was supported by University of Toronto Fellowship Award. J.R.S. is supported by a Catalyzing the Talent Pipeline Scholarship from the David Dime Family Catalyst Initiative in Molecular Genetics at the University of Toronto. J. L. was supported by Canada Graduate Scholarship, University of Toronto Fellowship and SickKids Restracomp Scholarship. Cadia Chan was supported by 02003PJT-437197 to MDW. This paper was generated with contributions from University of Toronto Master of Science theses from Katarina Maksimovic and Jooyun Lee.

## Author contributions

K.M., R.M., J.R.S., C.C., A.Z., J.L., H.K.Y. and J.P. conceived and designed the project. K.M., J.R.S., A.Z., J.L., O.B.G., M.M.M.Y., S.K., T.N., C.L., Y.F., M.N.M, J.Y. and C.S.K. performed the experiments, curated the data, analyzed and interpreted the results. R.M., C.C. and M.D. performed the bioinformatic analysis and interpreted the results. L.Y.W., J.Lefebvre., M.D.W., H.Y. and J.P. supervised, analyzed and interpreted the data. K.M., R.M., J.R.S., C.C., A.Z., J.L., H.K.Y. and J.P. wrote the first draft of the manuscript. All authors edited the manuscript, read and approved the final version.

## Data Availability Statement

Authors confirm that all relevant data are included in the paper and its Supplementary Information. Our cerebellar RNA seq data and motor neuron (ChAT)-specific RNA seq data have been deposited in Gene Expression Omnibus: https://www.ncbi.nlm.nih.gov/geo/query/acc.cgi?acc=GSE343867

## METHODS

### Mouse husbandry

All mouse procedures were performed under the approval of the Animal Care Committee at The Centre for Phenogenomics (TCP). Mice were kept on a 12-hour light/dark cycle with *ad libitum* access to food and water. Heterozygous (*Matr3^S85C/+^*) mice were crossed to generate wildtype (*Matr3^+/+^*), heterozygous (*Matr3^S85C/+^*), and homozygous (*Matr3^S85C/S85C^*) littermates. Mice were aged according to the timepoints described in each figure. Unless otherwise specified, both male and female mice were used without discrimination.

For TRAP-seq experiments, B6.FVB(Cg)-Tg(Chat-EGFP/Rpl10a,Slc18a3)DW167Htz/J (Jackson Laboratories #030250; Chat-EGFP/Rpl10a) mice were crossed to *Matr3^S85C/+^* mice, and their progeny were crossed to each other to generate wildtype (*Matr3^+/+^*) and homozygous (*Matr3^S85C/S85C^*) that were positive for the Chat-EGFP/Rpl10a transgene. Mice were aged and collected at denoted time points.

### Genotyping

Genomic DNA from mouse tissue was isolated as previously described^96^. DNA from ear or tail biopsies were genotyped using simple allele-discriminating PCR assay. For the MATR3 S85C mutation, the following primers were used: Common_Fwd 5′-GCGTTACCATTTTTGAAGCAA-3′; WT_Rev 5′-CCTCTACTTCCAATGTTAAATATGG-3′; S85C _Rev 5′-CCTCTACTTCCAATATTGAATATGC-3′. For the Chat-EGFP/Rpl10a transgene, the following primers were used: IntCtrl_Fwd 5′-AGTGGCCTCTTCCAGAAATG-3′; IntCtrl_Rev 5′-TGCGACTGTGTCTGATTTCC-3′; Tg_Fwd 5′-TCATAGAGGCGCAGAGTTCC-3′; Tg_Rev 5′-CTGAACTTGTGGCCGTTTAC-3′.

### Immunofluorescence

Mice that were 3 weeks or older were deeply anaesthetized using gaseous isoflurane then transcardially perfused using 1X phosphate-buffered saline (PBS) followed by 4% paraformaldehyde (PFA) in PBS. Mice that were less than 2 weeks old were anaesthetized using gaseous isoflurane and decapitated using sharp scissors. Brains, spinal columns, and tibialis anterior (TA) muscles were dissected and placed into 4% PFA for fixation. TA muscles were fixed for 10 minutes at room temperature (RT), washed with PBS three times for 5 minutes each, then stored at 4°C in PBS. Brains and spinal cords were fixed for 48 hours at RT, rocking gently, then washed with PBS 3 times for 10 minutes each, rocking gently. Fixed brains and spinal cords to be cryoembedded were first cryoprotected in 30% sucrose in PBS until saturated. For cerebellar sections, brains were embedded sagittally in paraffin and sectioned using a microtome (Leica RM2235) at 5 µm or embedded sagittally in optimal cutting temperature (OCT) compound (Tissue-Tek 4583) and sectioned using a cryostat (Thermo Scientific HM525) at 40 µm. For spinal cord sections, the spinal cord was removed from the spinal column, and the lower lumbar region (L4-L6) was embedded in paraffin or OCT. Spinal cords were transversely sectioned at 8 µm for paraffin-embedded samples and at 40 µm for cryoembedded samples.

Paraffin sections on positively-charged glass slides were rehydrated and washed 5 minutes for each step using xylene (3 times), 100% ethanol (2 times), 95% ethanol, 70% ethanol, water, PBS. Heat-induced epitope retrieval was performed with sodium citrate buffer (pH 6.0, in PBS) or Tris EDTA buffer (10 mM Tris, 1 mM EDTA, 0.05% Tween-20, pH 9). Sections were partitioned and permeabilized using 0.05% Tween-20 in PBS (PBST), then blocked for 1 hour at RT using 10% normal donkey serum and 0.25% Triton X-100 in PBST. Sections were then incubated with primary antibody diluted in blocking buffer overnight at 4°C. The following primary antibodies were used: MATR3 C-term (rabbit polyclonal, ab84422, 1:400), ChAT (goat polyclonal, ab144P or NBP1-30052, 1:200), NeuN (mouse monoclonal, MAB377, 1:200), calbindin (mouse monoclonal, Swant-300, 1:300), calbindin (goat polyclonal, AF3320, 1:200), phospho-c-JUN (Ser73) (rabbit monoclonal, 3270S, 1:200), NGFR (rabbit monoclonal, 8238, 1:500). Sections were washed with PBST three times, 5 minutes each, and incubated with secondary antibodies (488/555/647 Alexa Fluor donkey anti-rabbit/mouse/goat IgG (H + L), 1:500 from ThermoFisher Scientific) and DAPI (Millipore D6210, 1:1000) for 1 hour at RT. Sections were washed with PBST three times, 5 minutes each, and mounted with ProLong Gold Antifade mountant (ThermoFisher Scientific, P36930).

Cryoembedded tissue sections were put into PBS in a 12-well plate. All washes or incubations were performed on a rocker. Sections were permeabilized with 0.3% Triton X-100 in PBS (PBSTx) followed by 1 hour of blocking in 10% normal donkey serum in PBSTx. Overnight incubation of the sections at 4°C was performed in blocking buffer with primary antibodies diluted in concentrations as per the previous paragraph. Three 5-minute washes were done with PBSTx. Sections were incubated for 2 hours at RT in secondary antibodies diluted in blocking buffer. Another three 5-minute washes in PBSTx were completed before a 30-minute incubation with DAPI in PBSTx. Sections were washed a final three times for 5 minutes each. Sections were then mounted on neutral slides with ProLong Gold Antifade mountant.

All slides were imaged using SP8 Leica DM6000 confocal microscope (Leica Microsystems, Wetzlar, Germany), Nikon Eclipse Ti2E, or Leica Stellaris DMI8.

### Whole-mount immunostaining of tibialis anterior muscle

Tibialis anterior (TA) muscles collected from perfused mice were teased into thin pieces with membrane and fat removed. Muscle tissue was incubated in 0.1 M glycine in 1X PBS (pH 3.0) for 30 minutes at RT, rocking gently, to block free aldehydes. Muscle tissue was then permeabilized in 2% Triton X-100 in 1X PBS for 30 minutes at RT, rocking gently. Tissues were blocked in blocking buffer comprised of 4% Bovine Serum Albumin and 1% Triton X-100 in 1X PBS for 30 minutes at RT, rocking gently. Tissues were then incubated with primary antibody diluted in blocking buffer overnight at 4°C. The following primary antibodies were used: synaptophysin antibody (rabbit polyclonal, Synaptic Systems 101 002, 1:200), synapsin I antibody (rabbit polyclonal, ab64581, 1:200), and neurofilament H antibody (chicken polyclonal, ab4680, 1:200). Tissues were washed with PBS three times, 30 minutes each, then incubated in secondary antibodies (488 Alexa Fluor donkey anti-rabbit and 488 Alexa Fluor goat anti-chicken, 1:250 from ThermoFisher Scientific), α-bungarotoxin (Alexa Fluor 555 conjugate, Invitrogen B35451, 1:500), and DAPI (Millipore D6210, 1:1000) for 2 hours at RT, rocking gently. Tissues were washed with PBS three times, 10 minutes each, then mounted on concavity slides using ProLong Gold Antifade mountant (ThermoFisher Scientific, P36930). All slides were imaged using SP8 Leica confocal microscope (Leica Microsystems, Wetzlar, Germany).

### Motor neuron quantification

8 µm paraffin sections of transversely sectioned L4-L6 lumbar spinal cords were stained and imaged. A confocal z-series was imaged for each section at 40X magnification, 0.75 zoom, step size of 0.5 μm. Motor neurons were manually counted in the ventral horns based on ChAT and NeuN staining (α-motor neurons are positive for ChAT and NeuN; γ-motor neurons are positive for ChAT and negative for NeuN). Reduction of MATR3 staining in motor neurons with a visible nucleus (as seen using DAPI and NeuN) was manually counted based on staining levels relative to surrounding cells. 3 to 4 sections were averaged per animal.

### Purkinje cell quantification

5 µm paraffin sections of sagittally sectioned brains were stained and imaged. A confocal z-series was imaged for each section at 5X magnification, 0.75 zoom, step size of 0.5 μm. Representative images of the Purkinje cell layer were imaged at 63X magnification, step size of 0.5 μm. Purkinje cells were manually counted based on positive calbindin staining. MATR3 staining reduction was manually counted based on staining levels relative to surrounding cells. 4 sections were averaged per animal.

### Fluorescence intensity quantification

40 µm sagittal cryoembedded sections were stained and imaged. A z-series was imaged for each section with tile scanning (15% overlap) at 20X magnification, step size of 0.5 µm. Representative images of the Purkinje cell layer were imaged with tile scanning (15% overlap) at 63X magnification, step size of 0.5 µm. Cerebella were traced and integrated densities were measured using ImageJ. Values are given from the averages of 4 sections per animal.

### Neuromuscular junction denervation and endplate area quantification

Teased TA muscle was stained and imaged. About 20 to 30 confocal z-series were imaged per animal at 40X magnification, 0.75 zoom, step size of 3 µm. Over 100 neuromuscular junctions (NMJs) with connections to presynaptic axons were imaged per animal. The number of NMJs with presynaptic axonal swelling (blebbing) was manually counted. The number of NMJs with partial overlap (>1%, <80%) between the presynaptic terminal and post-synaptic terminal was manually counted (denervation). The endplates marked by α-bungarotoxin were manually outlined and their areas were quantified using ImageJ and averaged for each animal. One or both TA muscles were used for each animal, 5 animals per genotype.

### Statistical analysis – Purkinje cells, motor neurons, fluorescence intensity, neuromuscular junctions

Statistical significance was determined using unpaired, two-tailed Student’s *t*-test. All statistical tests were performed using GraphPad Prism 10 software package and data are displayed as mean ± SEM, with each datapoint representing an animal. P-values are as shown.

### Tissue collection and processing for RNA-sequencing

For bulk sequencing of the cerebellum, cerebella were collected from *Matr3^+/+^* and *Matr3^S85C/S85C^* mice at 2, 3, 4, and 6 weeks of age (n=3 animals/genotype; only females) and flash frozen in dry ice and stored at – 80 °C. For RNA extraction, the tissue samples were added in 1ml of Trizol reagent (Invitrogen, 15596018) and homogenized using tubes filled with Zirconium beads (OPS Diagnostics, PFAW 1400-100-19) and a tissue homogenizer, MagNA Lyser (Roche). Samples were then incubated at room temperate (RT) for 10 mins and mixed with chloroform (Sigma-Aldrich, 1731042) and centrifuged. The supernatant was transferred to a new tube, and RNA was precipitated using isopropanol. After centrifugation, the RNA pellet was washed with 75% ethanol and dried for 20 mins. After drying, the RNA pellet was resuspended in Diethyl pyrocarbonate DEPC-treated water (Ambion, AM9920) and incubated for 15 mins at 55°C. Samples were placed on ice for 30 mins before the concentration was measured using the Nanodrop (ThermoFisher). Total RNA samples were then stored at -80°C before sent to Azenta Life Sciences, Inc. for mRNA library preparation, which were paired end (150 bp in length) sequenced on the HiSeq 2 flow cell.

For motor neuron TRAP-seq, 2 to 3 spinal cords from female *Matr3^+/+^* and *Matr3^S85C/S85C^* mice that were positive for the Chat-EGFP/Rpl10a transgene were pooled per replicate. Spinal cords were homogenized in 1 mL of dissection buffer; otherwise, bead preparation, tissue homogenization, and immunopurification (IP) was performed as previously described^74, 75, 97^. RNA was extracted from washed beads using the Qiagen RNeasy Mini kit (Qiagen 74104). An RNase-free DNase digestion step was performed (Qiagen 79254). Extracted RNA yield was quantified using a Nanodrop (ThermoFisher). IPed RNA samples were then stored at -80°C before sent to SickKids Centre for Applied Genomics (TCAG) core for mRNA library preparation, which were then paired end (150 bp in length) sequenced on the Novaseq S4 flowcell.

### RNA-seq reads processing and analysis

Raw sequencing reads were assessed for quality using FastQC v0.11.9^98^, evaluating per-base quality scores, GC content, and adapter contamination. Adapter trimming and low-quality read filtering were performed with fastp v0.20.0^99^, and processed reads were aligned to the mouse reference genome GRCm38 (GENCODE release v23) using STAR v2.7.9a^100^. The STAR alignment index was constructed from FASTA sequences and GTF annotation files obtained from the GENCODE portal.

Gene-level read counts were quantified from STAR alignments using the --quantMode GeneCounts parameter, and differential expression analysis was performed using DESeq2^101^ following median-of-ratios normalization. Genes with a mean read count below 100 across all samples were excluded prior to testing. Differentially expressed genes (DEGs) were defined by an adjusted p-value (padj) of 0.05 or below combined with an absolute log2 fold change of 0.263 or greater, corresponding to a minimum 20% change in expression^102^. Volcano plots for the cerebellum time course (Weeks 2, 3, 4, and 6) and motor neuron stages (Weeks 4 to 6, 12 to 15, and 30 to 33) were generated in R.

Temporal expression dynamics were visualized using heatmaps for both tissue types. For the cerebellum, DEGs identified at Week 6 were tracked across all time points by extracting log2FC values at each week and displaying them as longitudinal trajectories (Figure 4E). For motor neuron gene expression, raw counts were first batch corrected using the ComBat_seq function (sva R package) to eliminate technical variation. In the motor neuron heatmap, DEGs from Weeks 12 to 15 served as the reference gene set. A baseline mean expression was calculated from wild-type samples at Weeks 4 to 6, and individual counts across all time points were normalized to this baseline to derive relative fold changes, shown in Figure 6E. In both the cerebellum and motor neurons, continuous trajectory profiles were generated for candidate genes using the ggplot2 R package. Longitudinal expression trends between *Matr3^+/+^* and *Matr3^S85C/S85C^* groups were visualized by applying local polynomial regression fitting (loess) to individual biological replicates and summarized group averages (log_10_ scaled normalized counts) across timepoints.

To further evaluate coordinated behavior of Integrated Stress Response (curated ATF4 target gene set^89, 103, 104^) and Inflammatory Response, targeted Gene Set Enrichment Analysis (GSEA) was performed using clusterProfiler (v4.12) and GSEA was run across ranked gene lists with a minimum gene set size of 5. Enrichment trajectories and barcode plots were rendered using the enrichplot and GseaVis packages.

Pathway enrichment analysis was performed using Enrichr^105^. Cross-model comparison was performed using differentially expressed genes identified in Shadrach et al., 2021^13^ and Sun et al., 2015^12^.

### Single cell mapping using published single-nuclei RNA-seq

Single nuclei RNA-seq (snRNA-seq) was performed by Kozavera et al., 2021^60^ in 60-day-old adult female and male mouse cerebellar cortex. The annotated adult mouse Seurat object was obtained from Single Cell Portal (SCP795) https://singlecell.broadinstitute.org/single_cell/study/SCP795/. As the Seurat object was generated using Seurat v2 in the original study, UpdateSeuratObject() was first used to ensure compatibility with Seurat v3.1.4, which we used in this study. Gene markers for each cell type was identified using Seurat FindMarkers() with test.use = “wilcox”, and default parameters (logfc.threshold = 0.25, min.pct = 0.1). Genes with avg_log2FC > 0.25 and p_val_adj < 0.05 were considered positive gene markers for each cell type. If a gene met these criteria for multiple cell types, it was assigned to the cell type with the highest percentage of cells expressing the given gene. To evaluate the expression of cell-type specific gene markers in our study, we overlapped bulk DE gene lists we identified in this study and our previously published bulk RNA-seq study with these positive cell-type gene markers identified from Kozavera et al., 2021^60^.

### cDNA synthesis and qRT-PCR

1-2 micrograms of total RNA extracted from each tissue sample were used to generate cDNA using random hexamer primers and the M-MLV reverse transcriptase (Invitrogen) according to the manufacturer’s direction. For confirmation of enrichment following TRAP, 120 ng of RNA from the input, IP, and flowthrough samples were used to generate cDNA as above. qRT-PCR was performed with cDNA using iTAQ universal SYBR green Super Master Mix (Biorad Labs, 1725271) with readout using the Applied Biosystems ViiA 7 real-time PCR system with standard cycling parameters followed by standard melt curve analysis. Primers used for qRT-PCR are listed in a separate table (**Supplementary Table 1**).

ΔΔC_t_ was calculated with the housekeeping gene *Gapdh* or *36b4 (Rplp0)* as reference and wildtype samples as the calibrator. The relative expression or fold change was calculated using the 2^-ΔΔCt^ method. Statistical analysis was performed using GraphPad Prism and significance was calculated using unpaired t-tests.

### Protein extraction, Immunoblotting, and ELISA

For immunoblotting, samples were homogenized in RIPA buffer supplemented with protease and phosphatase inhibitors using a 5 mm stainless steel bead and a TissueLyser II, followed by sonication and clarification by centrifugation. Protein concentrations were determined using a BCA assay. Equal amounts of proteins were electrophoresed on a hand-casted bis-acrylamide gel in Tris-Glycine-SDS Running Buffer (25 mM Tris, 192 mM glycine, 0.1% SDS) then transferred onto 0.45 μM nitrocellulose membrane (Amersham Protran Premium) in Towbin’s transfer buffer (25 mM Tris, 192 mM Glycine, 20% Methanol), prior to blocking and probing with primary antibody diluted in blocking buffer (5% BSA in TBS). Blots were probed using the following primary and secondary antibodies: mouse monoclonal anti-Vinculin (Sigma Aldrich, V9131, 1:1000), rabbit monoclonal anti-NGFR (Cell Signaling Technology, 8238, 1:1000), IRDye680RD conjugated goat polyclonal anti-mouse IgG (Licor, 925-68070, 1:7500) antibody, and IRDye 800CW conjugated goat polyclonal anti-rabbit IgG (Licor, 926-32211, 1:7500) antibody. Blots were imaged on the Li-Cor Odyssey Fc imager. Band density was quantified using *ImageJ*.

For the FGF21 ELISA, samples were homogenized in PBS supplemented with protease inhibitor using a 5 mm stainless steel bead and a TissueLyser II, followed by sonication, one freeze-thaw cycle, and clarification by centrifugation at 4°C. Protein concentrations were determined using a BCA assay. The ELISA was performed as per manufacturer’s instructions (R&D Systems, MF2100). The concentrations determined from the standard curve were then normalized to the protein concentration and reported as pg of FGF21 per mg of proteins.

