## Supplementary Figures for "Early MATR3 loss and distinct neurodegenerative molecular signatures precede the onset of neuropathology in motor neurons and Purkinje cells of MATR3 S85C knock-in mouse model of ALS"

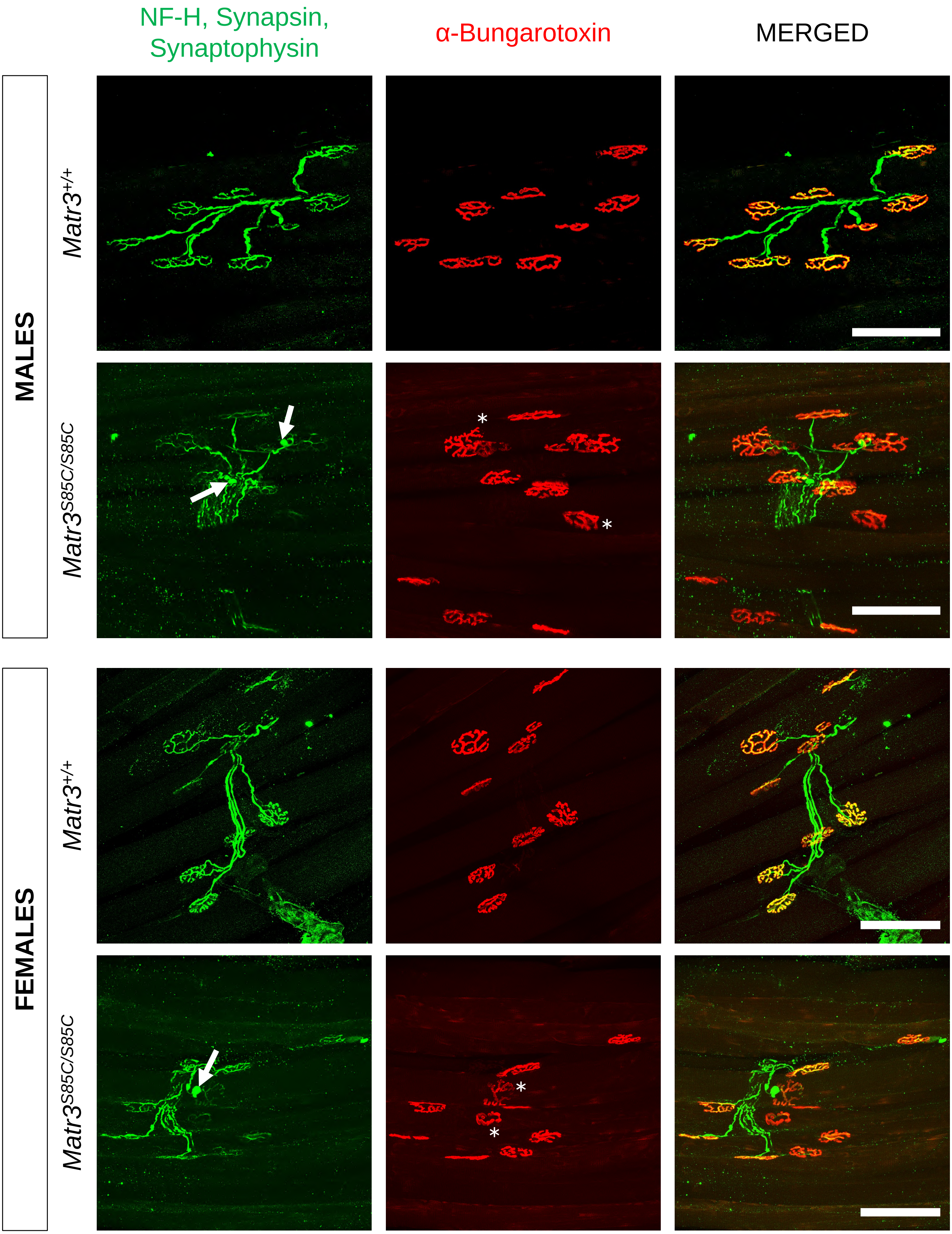
NF-H, Synapsin,
Synaptophysin
α-Bungarotoxin
MERGED
MALES
FEMALES
Matr3+/+
Matr3S85C/S85C
Matr3+/+
Matr3S85C/S85C

**Supplementary Figure 1. Neuromuscular junction pathology is evident in male and female *Matr3*$^{S85C/S85C}$ mice at 30 weeks of age.**

Representative images of the presynaptic axons (Neurofilament H, Synapsin, Synaptophysin; green) and postsynaptic motor endplates (alpha-bungarotoxin; red) of the neuromuscular junctions (NMJs) in the tibialis anterior muscle of male and female *Matr3*$^{+/+}$ and *Matr3*$^{S85C/S85C}$ mice at 30 weeks of age. Scale bars indicate 100 µm. White arrows point to presynaptic axonal blebbing and asterisks mark NMJs displaying partial denervation.

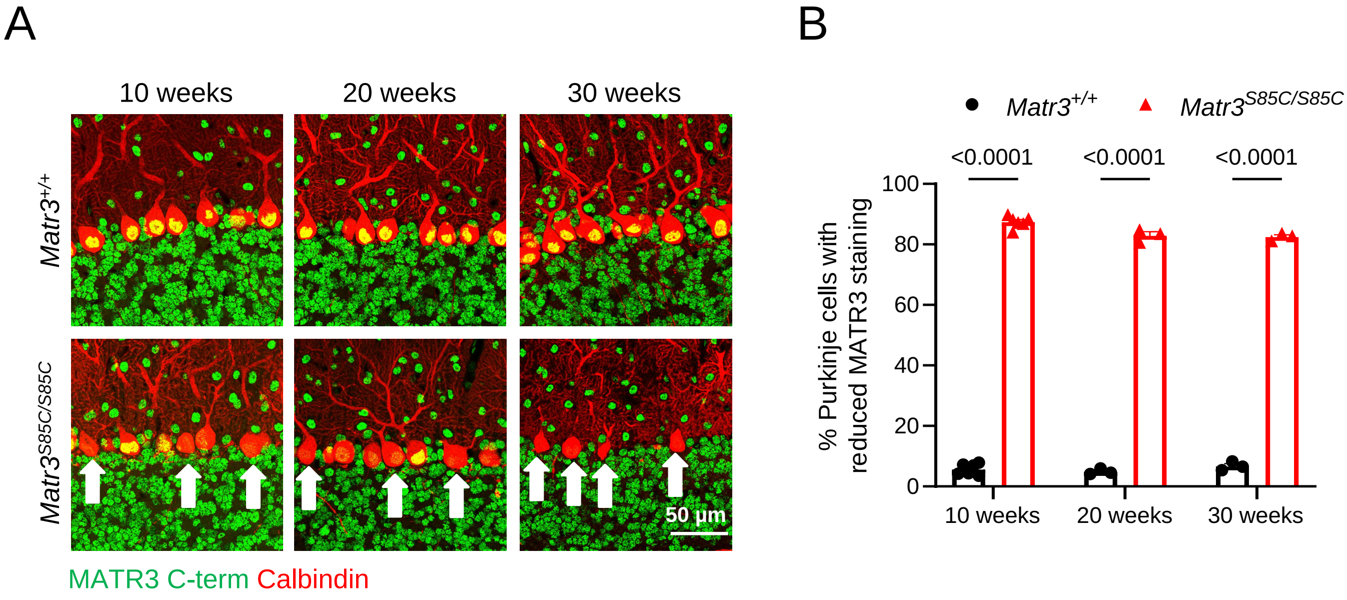


**Supplementary Figure 2. Reduced MATR3 staining in Purkinje cells of *Matr3*$^{S85C/S85C}$ mice at 10, 20, and 30 weeks of age.**

**(A)** Representative images of MATR3 and Calbindin immunostaining in the cerebellum of male and female *Matr3*$^{+/+}$ and *Matr3*$^{S85C/S85C}$ mice at the indicated ages. Scale bars indicate 50 µm. Representative Purkinje cells with reduced MATR3 signal intensity are marked by white arrows. N=3-4 mice per sex per genotype, bar heights depict mean ± SEM with each dot representing the average value quantified from four sections per single animal, statistical significance ($p<0.05$) determined by multiple unpaired t-tests, p-values as shown.

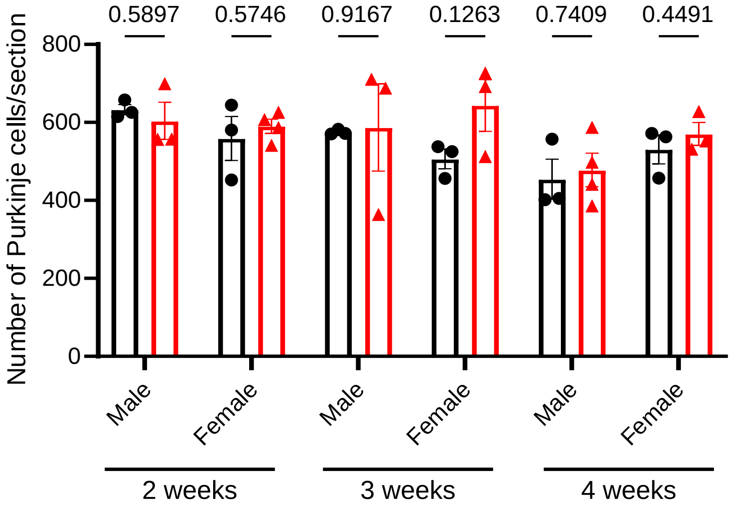


**Supplementary Figure 3. No difference in the number of Purkinje cells in cerebellar sections between *Matr3*$^{+/+}$ and *Matr3*$^{S85C/S85C}$ male and female mice at 2, 3, and 4 weeks of age*.***

Quantification of the total number of Purkinje cell soma throughout the whole cerebellum. N=3-4 mice per sex per genotype, bar heights depict mean ± SEM with each dot representing the average value quantified from four sections per single animal, statistical significance ($p<0.05$) determined by multiple unpaired t-tests, p-values as shown.

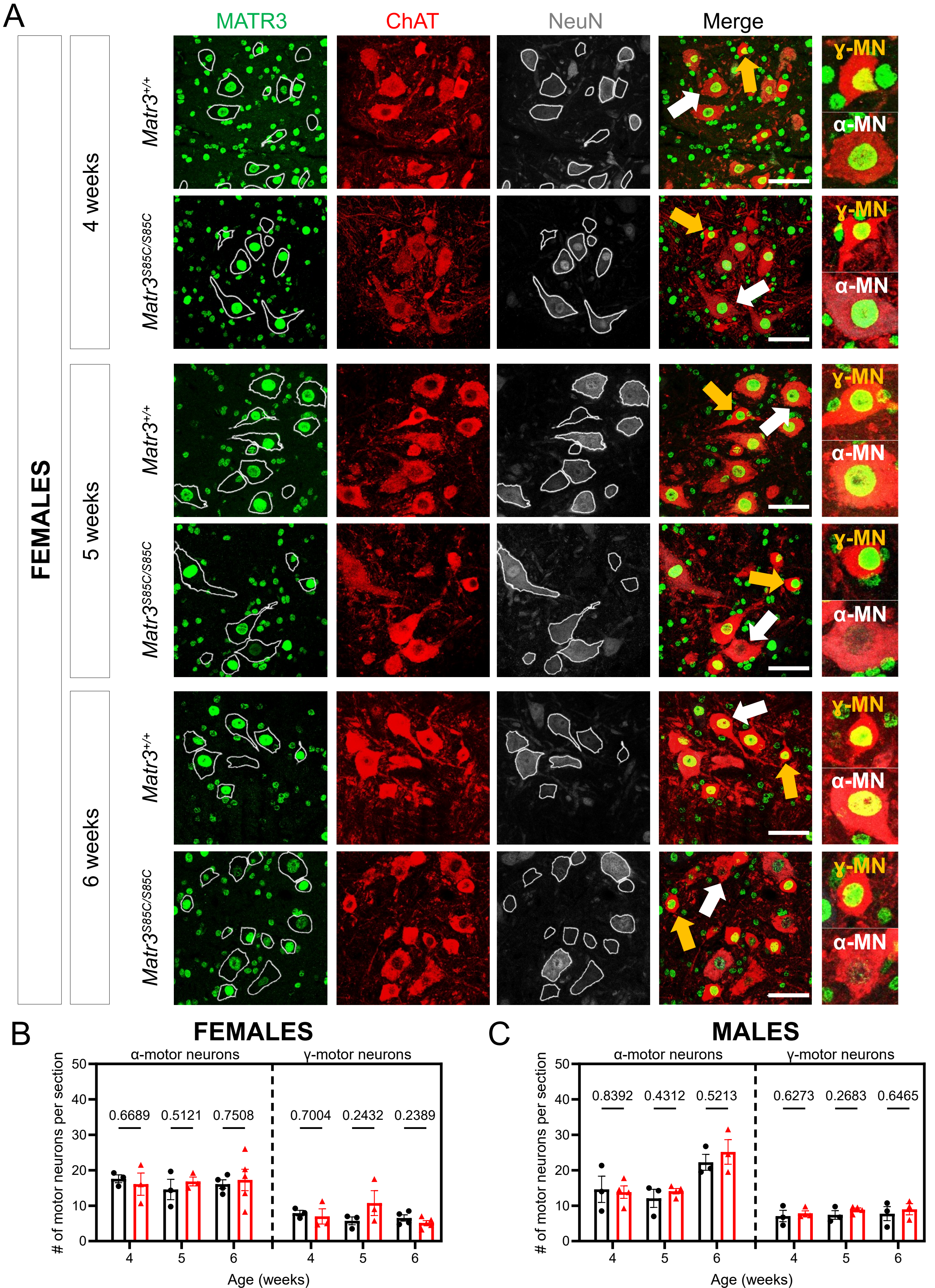

A
MATR3
ChAT
NeuN
Merge
FEMALES
4 weeks
5 weeks
6 weeks
Matr3+/+
Matr3S85C/S85C
γ-MN
α-MN
B
FEMALES
α-motor neurons
γ-motor neurons
### of motor neurons per section
0.6689
0.5121
0.7508
0.7004
0.2432
0.2389
Age (weeks)
C
MALES
0.8392
0.4312
0.5213
0.6273
0.2683
0.6465

**Supplementary Figure 4. Reduced MATR3 staining in the α-motor neurons but not in the γ-motor neurons of female *Matr3*$^{S85C/S85C}$ mice occurs before the onset of motor dysfunction and loss of motor neuron soma.**

**(A)** Representative images of MATR3, ChAT, and NeuN immunostaining in the α-motor neurons (α-MN; $ChAT^{+}NeuN^{+}$) and γ-motor neurons (γ-MN; $ChAT^{+}NeuN^{-}$) in the lumbar spinal cord of female *Matr3*$^{+/+}$ and *Matr3*$^{S85C/S85C}$ mice at the indicated ages. Scale bars indicate 50 µm. Representative motor neurons are magnified in the inset and are indicated with white (α-MN) or orange (γ-MN) arrows.

**(B-C)** Quantification of the total number of α-MN or γ-MN throughout the ventral horn of the lumbar spinal cord for female and male mice. N=3-5 mice per sex per genotype, bar heights depict mean ± SEM with each dot representing the average value quantified from four sections per single animal, statistical significance ($p<0.05$) determined by multiple unpaired t-tests, p-values as shown.

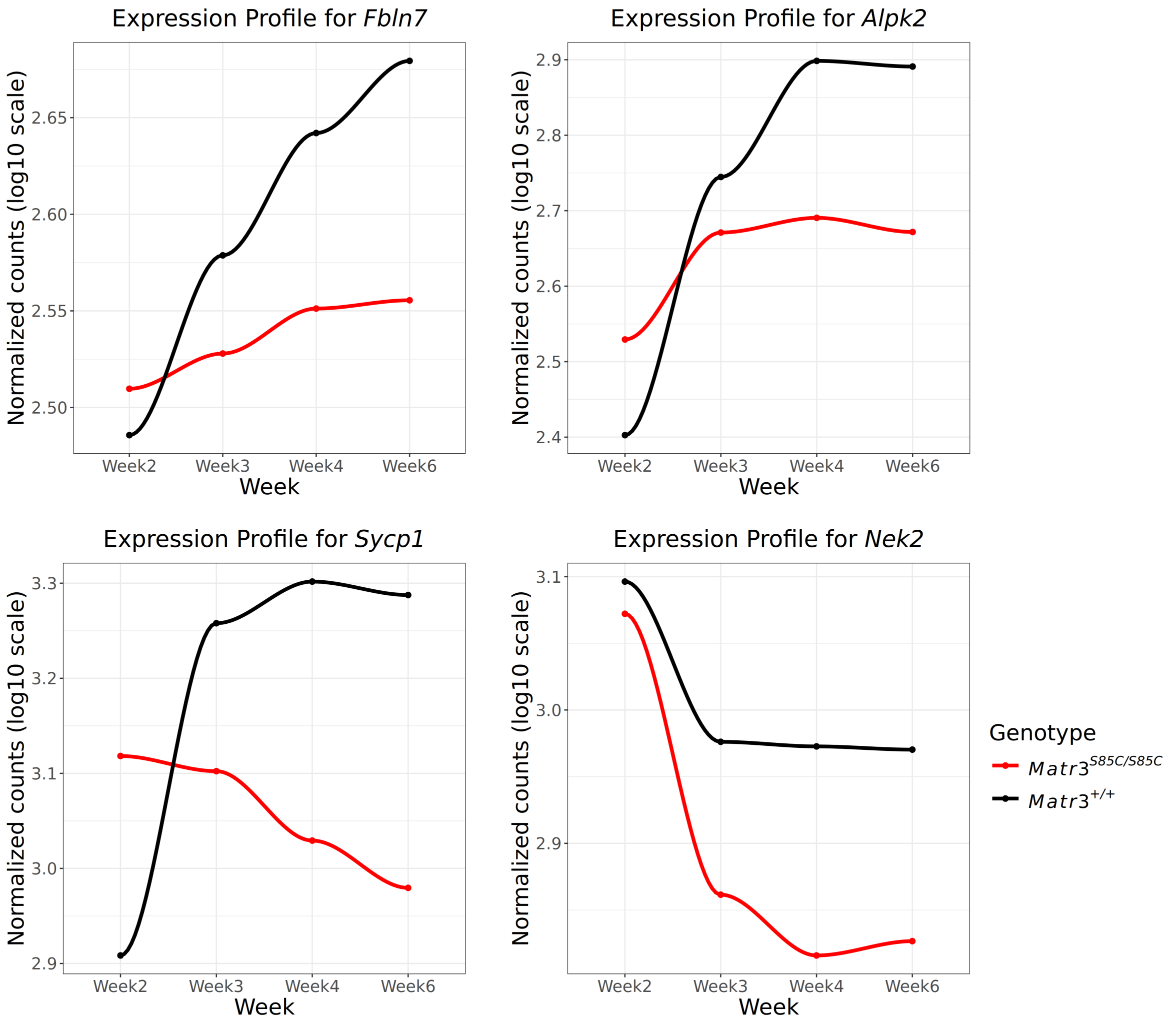


**Supplementary Figure 5.** Temporal expression profiles of selected genes from bulk cerebellar RNA-sequencing at Weeks 2, 3, 4, and 6. Lines represent the mean normalized expression (log10 counts) for each genotype.

6-week-old cerebellum

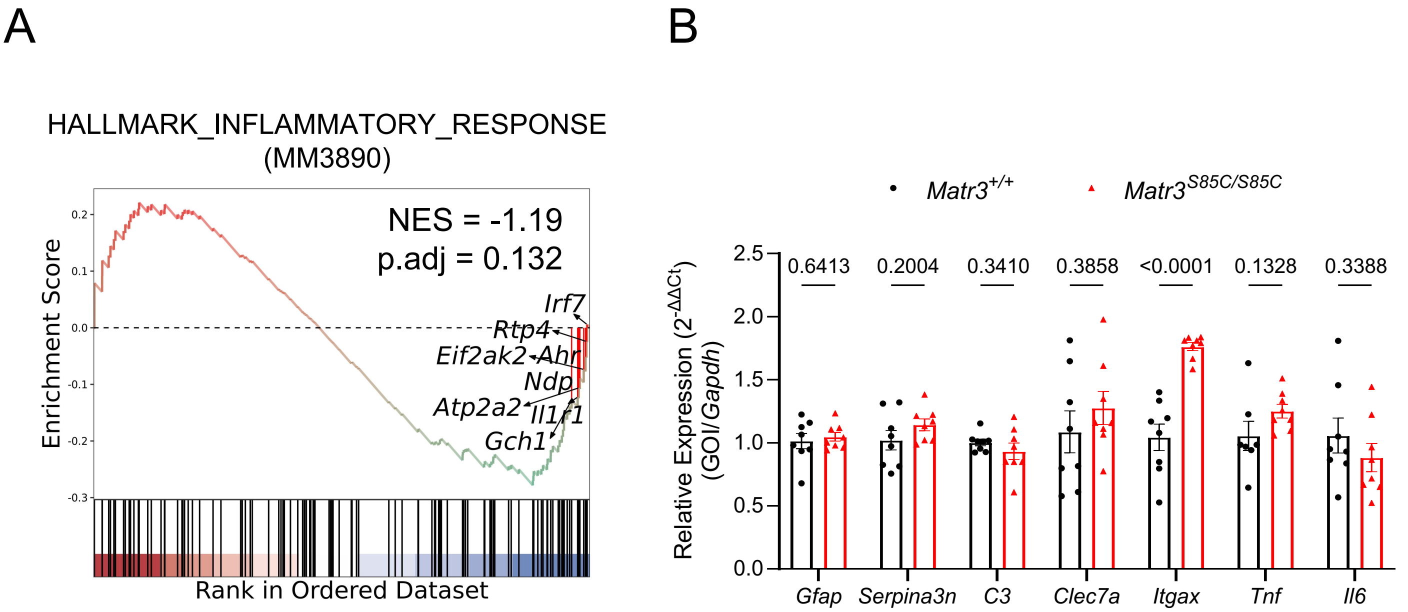


**Supplementary Figure 6. Transcriptomic analyses reveal minimal evidence of inflammatory pathway activation in the cerebellum of 6-week-old *$Matr3^{S85C/S85C}$* mice.**

**(A)** Gene Set Enrichment Analysis for genes associated with the inflammatory response (MM3890 Hallmark gene set) in 6-week cerebellum revealed a modest but not statistically significant negative enrichment.

**(B)** Relative expression ($\Delta\Delta C_t$ analysis) of inflammation-associated genes from total RNA isolated from the cerebellum of *$Matr3^{+/+}$* and *$Matr3^{S85C/S85C}$* mice at 6 weeks of age as measured by quantitative RT-PCR (N=8 mice per genotype, bar heights depict mean ± SEM with each dot representing a single animal, significance determined by multiple unpaired t-test).

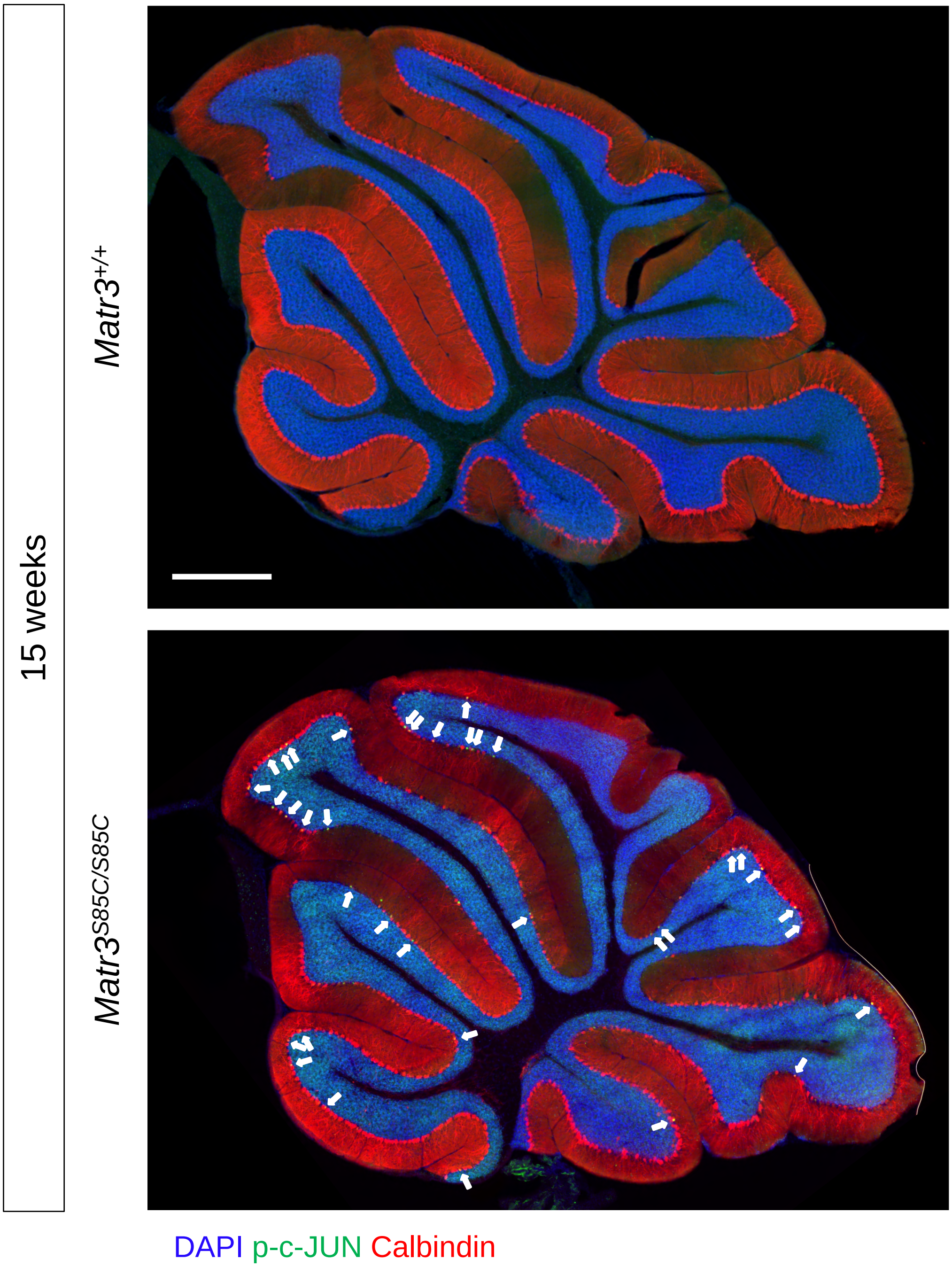


**Supplementary Figure 7. Whole cerebellum images corresponding to representative 15-week images in Figure 5F showing p-c-JUN staining.**

Whole cerebellum images of calbindin-positive Purkinje cells with nuclear phospho-c-JUN (p-c-JUN) staining at 15 weeks of age. Purkinje cells with nuclear p-c-JUN staining are indicated by a white arrow. Scale bar indicates 500 µm.

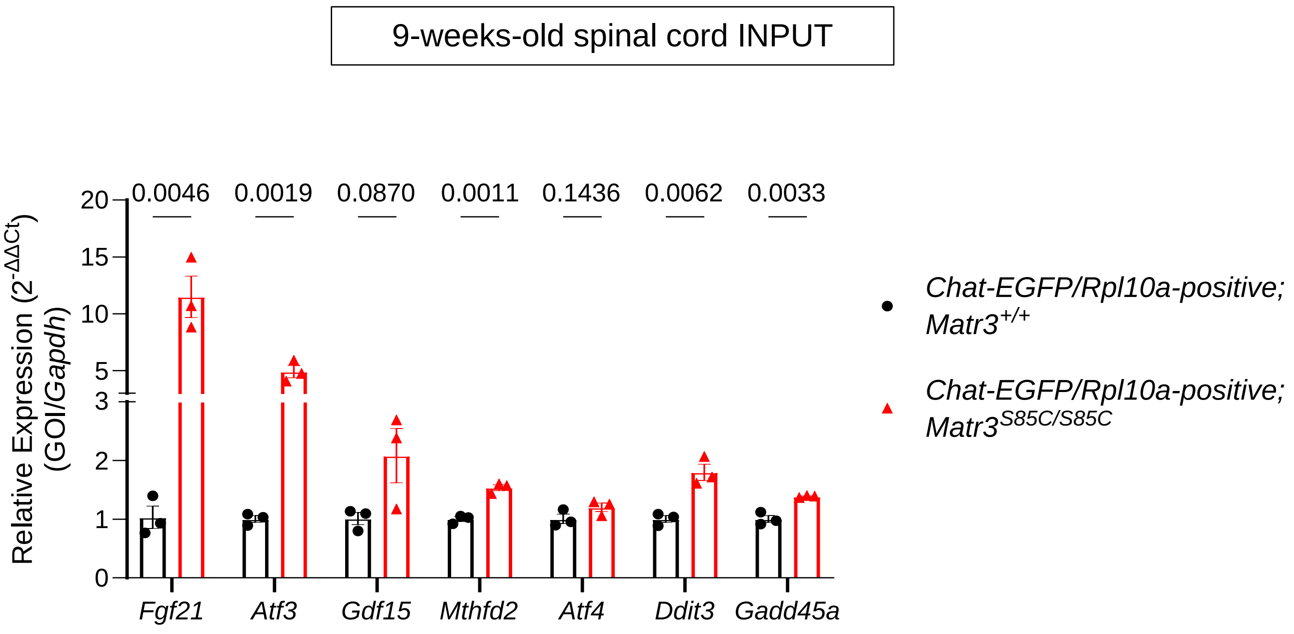


**Supplementary Figure 8. ISR-associated genes are significantly upregulated in 9-week-old spinal cord samples from *Chat-EGFP/Rpl10a-positive; Matr3$^{S85C/S85C}$* mice.**

Relative expression ($\Delta\Delta C_t$ analysis) of ISR-associated genes from total RNA isolated from the spinal cord of *Chat-EGFP/Rpl10a-positive; Matr3$^{+/+}$* and *Chat-EGFP/Rpl10a-positive; Matr3$^{S85C/S85C}$* mice at 9 weeks of age as measured by quantitative RT-PCR (N=3 mice per genotype, bar heights depict mean ± SEM with each dot representing a single animal, significance determined by multiple unpaired t-test).

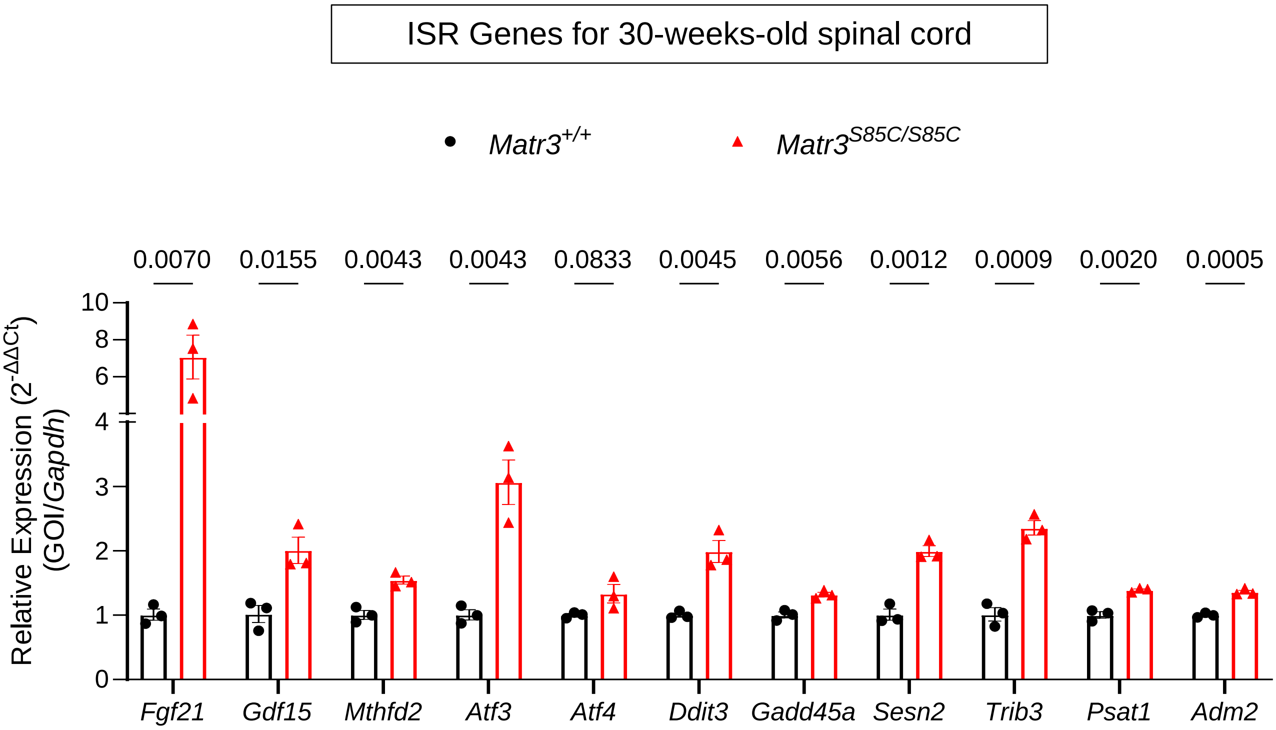


**Supplementary Figure 9. ISR-associated genes are significantly upregulated in 30-week-old spinal cord samples from *Matr3*$^{S85C/S85C}$ mice.**

Relative expression ($\Delta\Delta C_t$ analysis) of ISR-associated genes from total RNA extracted from the spinal cord of *Matr3*$^{+/+}$ and *Matr3*$^{S85C/S85C}$ mice at 30 weeks of age as measured by quantitative RT-PCR (N=3 mice per genotype, bar heights depict mean ± SEM with each dot representing a single animal, significance determined by multiple unpaired t-test).

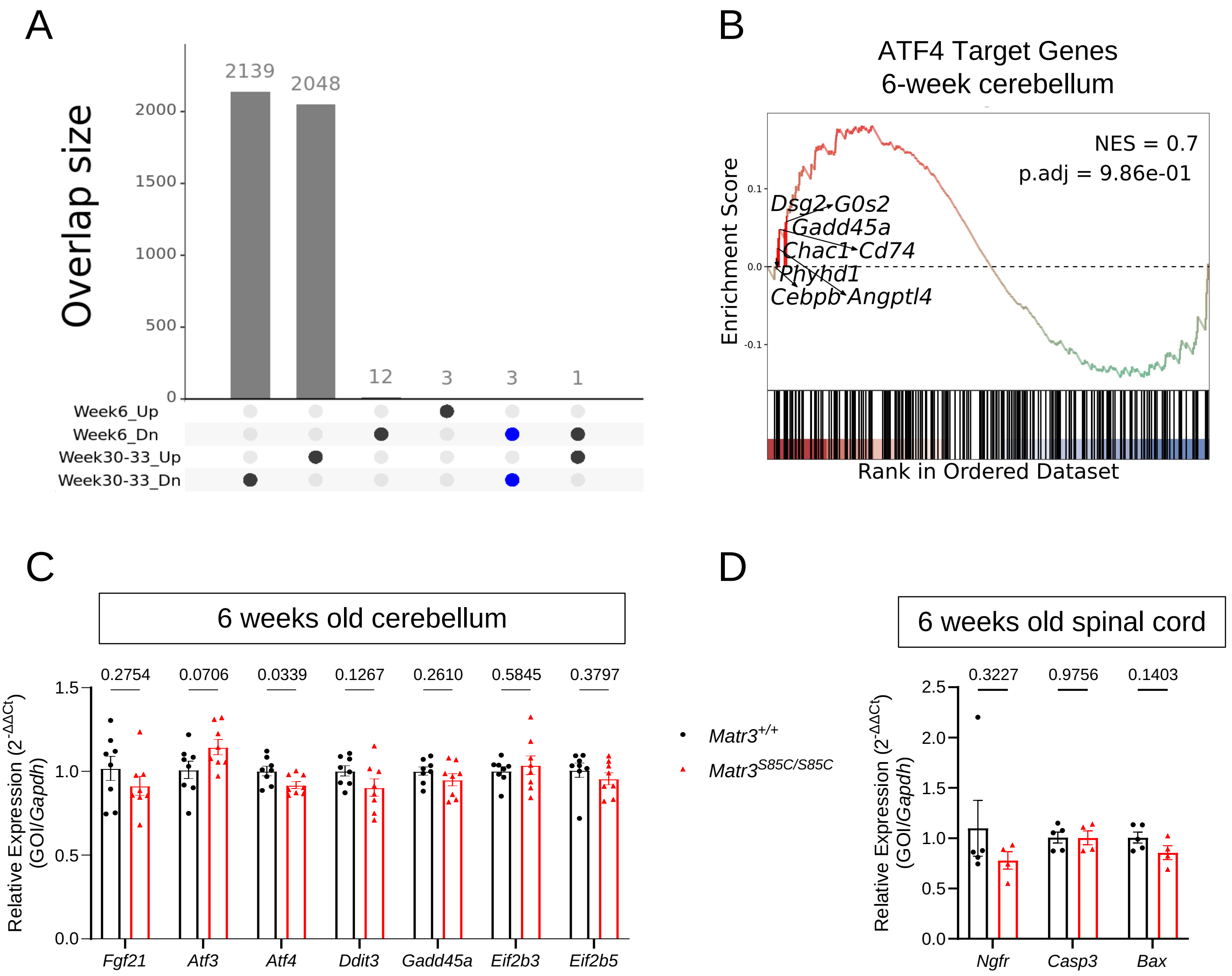


**Supplementary Figure 10. Few differentially expressed genes common in both cerebellar and motor neuron RNA seq.**

**(A)** UpSet plot showing the overlap of differentially expressed genes between the 6-week cerebellar RNA-seq dataset and the 30–33-week motor neuron RNA-seq dataset. Only three genes (*Nell1*, *Rbm3*, and *Upt18*) were commonly downregulated in both datasets.

**(B)** Gene Set Enrichment Analysis of ATF4 Target genes in 6-week cerebellum showed no significant enrichment.

**(C-D)** Relative expression ($\Delta\Delta C_t$ analysis) of ISR-associated or NGFR-JNK-associated genes from total RNA extracted from the cerebellum or the spinal cord of *Matr3*$^{+/+}$ and *Matr3*$^{S85C/S85C}$ mice at 6 weeks of age as measured by quantitative RT-PCR (N=4-8 mice per genotype, bar heights depict mean ± SEM with each dot representing a single animal, significance determined by multiple unpaired t-test).

**Supplementary Table 1. Sequences of primers used for the quantitative real time PCR (qRT-PCR).**

| Gene | Forward Primer (5' → 3') | Reverse Primer (5' → 3') |
|---|---|---|
| *Adm2* | CAGACAACAGACGCAGCCCA | GAAGGAATCTTAGCTGGGGGC |
| *Aldh1l1* | GCAGGTACTTCTGGGTTGCT | GGAAGGCACCCAAGGTCAAA |
| *Alpk2* | AGCCTGTGAACTGAGTCTCCT | CTTGACCCACTGCCCACATTT |
| *Atf3* | AGCGAAGACTGGAGCAAAATG | ACAAAGGGTGTCAGGTTAGCA |
| *Atf4* | GGCGTATTAGAGGCAGCAGT | GGATTTCGTGAAGAGCGCCA |
| *Bax* | GGAGCAGCTTGGGAGCG | AAAAGGCCCCTGTCTTCATGA |
| *Bcl2* | GAGTTCGGTGGGGTCATGTG | GCATGCTGGGGCCATATAGTT |
| *C3* | GCAGGTCATCAAGTCAGGCT | GGCTTTTCTCCCCAGAGGTC |
| *Casp3* | GAGCTTGGAACGGTACGCTA | GAGTCCACTGACTTGCTCCC |
| *Chat* | GACCAGCTAAGGTTTGCAGC | CAGGAAGCCGGTATGATGAGA |
| *Chrna1* | GAGTGGGTGCGGAAGGTTTT | CTGGATGGTCTTTTCATTGTGGA |
| *Chrng* | CTTCCAATCCCAGACTTACAGC | ACTCCCCATTCTCTGTGAAAG |
| *Clec7a* | AACGTCGTGACCCAAGCTAC | AAGGGCAGCACCTTTGTCAT |
| *Cx3cr1* | CGTGAGACTGGGTGAGTGAC | GGACATGGTGAGGTCCTGAG |
| *Ddit3* | GAGCTGGAAGCCTGGTATGA | GGTTTTTGATTCTTCCTCTTCGT |
| *Dner* | TGTGCCGTAGCGTGGG | AGTTCGCAGTGTGTTCCTTTA |
| *Eif2b3* | GCCAACAGACAGGTCCCCAA | CAGGATGAGCCAATGACAGAGT |
| *Eif2b5* | CCTGGAGGAACACAGGTTAAGA | GGCTGGGTGACGACTCTTTG |
| *Fasl* | TCCGTGAGTTCACCAACCAA | TGAGTGGGGGTTCCCTGTTA |
| *Fbln7* | AGCGTGGTGTGTCTTGCTAA | CCCTCCATTGTGACAAGGCT |
| *Fgf21* | GGGTCTACCAAGCATACCCC | CTGATCTCCAGGTGGGCTTC |
| *Gadd34* | CCTCTAAAAGCTCGGAAGGTACAC | ATCTCGTGCAAACTGCTCCC |
| *Gadd45a* | CAGAGCAGAAGACCGAAAGGATG | CAGAACGCACGGATGAGGGT |
| *Gapdh* | AGGTCGGTGTGAACGGATTTG | TGTAGACCATGTAGTTGAGGTCA |
| *Garnl3* | GAAGATTCCGAGTGGAGAATGG | TGTGGTACTCCGGGTTTTCTAA |
| *Gdf15* | GTCCCAACTCAACGCCGAC | GGACCCCAATCTCACCTCTGG |
| *Gfap* | GACCAGCTTACGGCCAACA | TTCATCTTGGAGCTTCTGCCT |
| *Il6* | TCTATACCACTTCACAAGTCGGA | GAATTGCCATTGCACAACTCTTT |
| *Itgax* | GAGGCTGCAAGCATCATTCG | GCATCAAAGTTCTCCACGCTG |
| *Mcl1* | AACGGGACTGGCTTGTCAAA | CTGATGCCGCCTTCTAGGTC |
| *Mnx1* | CCAAGATGCCGGACTTCAGC | TGGAACCAAATCTTCACCTGAGTCT |
| *Mthfd2* | AAGAATGTGGTAGTGGCTGGC | GACTGTGGCATCACCCCCG |
| *Nek2* | TTCCATCCTCAGCCATGAAGA | CCTGCACTTGGACTTGGCAA |
| *Ngfr* | CTAGGGGTGTCCTTTGGAGGT | CAGGGTTCACACACGGTCT |
| *Psat1* | TTGAAGGGGCACAGGTCAGT | CAGAGCGTGGCTGGTCAT |
| *Serpina3* | ATTTGTCCCAATGTCTGCGAA | TGGCTATCTTGGCTATAAAGGGG |
| *Sesn2* | CGAGACCCGAGAGGAGCATC | TGTAGACCCATCACCACCGC |
| *Sycp1* | ACTGCTGATCTCTTGAGCCTG | AAGGAGTCTGTTTGGGGGTT |
| *Tnf* | CTTCTCATTCCTGCTTGTGGC | ACTTGGTGGTTTGCTACGACG |
| *Trib3* | GCAGCACTTTAGCAGCGGA | GAGGTGTAGCTCGCATCTTGT |
| *Xbp1* spliced | TGCTGAGTCCGCAGCAGGTG | TGCCCAAAAGGATATCAGACTCA |
| *Xbp1* unspliced | CTCAGACTATGTGCACCTCTGC | TGCCCAAAAGGATATCAGACTCA |
| *Xiap* | TGCTGGACTGGGCTCCCTAT | TAGGACTTGTCCACCTTTTCTTAGG |
